# Processivity and Velocity for Motors Stepping on Periodic Tracks

**DOI:** 10.1101/684696

**Authors:** M.L. Mugnai, M.A. Caporizzo, Y.E. Goldman, D. Thirumalai

**Affiliations:** Department of Chemistry, The University of Texas at Austin, Austin, Texas; Department of Physiology, Pennsylvania Muscle Institute, Perelman School of Medicine, University of Pennsylvania, Philadelphia, Pennsylvania

## Abstract

Processive molecular motors enable cargo transportation by assembling into dimers capable of taking several consecutive steps along a cytoskeletal filament. In the well-accepted hand-over-hand stepping mechanism the trailing motor detaches from the track and binds the filament again in leading position. This requires fuel consumption in the form of ATP hydrolysis, and coordination of the catalytic cycles between the leading and the trailing heads. However, alternative stepping mechanisms exist, including inchworm-like movements, backward steps, and foot stomps. Whether all of these pathways are coupled to ATP hydrolysis remains to be determined. Here, in order to establish the principles governing the dynamics of processive movement, we present a theoretical framework which includes all of the alternative stepping mechanisms. Our theory bridges the gap between the elemental rates describing the biochemical and structural transitions in each head, and the experimentally measurable quantities, such as velocity, processivity, and probability of backward stepping. Our results, obtained under the assumption that the track is periodic and infinite, provide expressions which hold regardless of the topology of the network connecting the intermediate states, and are therefore capable of describing the function of any molecular motor. We apply the theory to myosin VI, a motor that takes frequent backward steps, and moves forward with a combination of hand-over-hand and inchworm-like steps. Our model reproduces quantitatively various observables of myosin VI motility measured experimentally from two groups. The theory is used to predict the gating mechanism, the pathway for backward stepping, and the energy consumption as a function of ATP concentration.

**Significance Statement:** Molecular motors harness the energy released by ATP hydrolysis to transport cargo along cytoskeletal filaments. The two identical heads in the motor step alternatively on the polar track by communicating with each other. Our goal is to elucidate how the coordination between the two heads emerges from the catalytic cycles. To do so, we created a theoretical framework that allows us to relate the measurable features of motility, such as motor velocity, with the biochemical rates in the leading and trailing heads, thereby connecting biochemical activity and motility. We illustrate the efficacy of the theory by analyzing experimental data for myosin VI, which takes frequent backward steps, and moves forward by a hand-over-hand and inchworm-like steps.

## Introduction

Long distance transport in cells is essential for the targeted delivery and recycling of sub-cellular machinery. Cytoskeletal filaments, actin and microtubules, supply the analogous roads and highways for the molecular motor families myosin, kinesin, and dynein that catalyze intracellular transport (1). In many examples of intracellular transport, only a few motors of a particular type are attached to the cargo (2–6), so they usually operate in very small groups. Dimeric processive molecular motors with two identical heads, such as myosin V (7), VI (8), and X (9), kinesin-1 (10), and cytoplasmic dynein (11) take multiple consecutive steps along their track for each diffusional encounter with the filament, enabling long-distance movement. During each step, the motor completes a catalytic cycle, in which it occupies a number of intermediate biochemical states. Thermodynamics imposes that each step be reversible, hence the cycle could be completed “backwards” as well, although coupling to the hydrolysis of ATP to ADP and inorganic phosphate (P_i_) guarantees that the motor progresses predominantly in one direction. Run termination occurs via stochastic dissociation of both the heads from the filament. Thus, long-distance movement is achieved by a strong association between the filament and the predominantly occupied biochemical intermediate states of the reaction cycle (high duty ratio) (12–15).

The movement of processive motors can be monitored using a variety of single-molecule measurements, for instance optical tweezers (16), fluorescence of bound probes (17), or light scattering from a small refractile particle (18), which are each suited to extract specific information over limited spatial and temporal ranges. Actin and microtubules are both polarized, and the specific motors move mainly towards one of the two filament ends. The predominant isoforms of all three motor families involved in long-distance transport form dimers having two actin-or microtubule-binding domains which bind successively in positions ahead of (the leading head, LH) or behind (the trailing head, TH) the center of mass. Determining the stepping mechanism, such as canonical hand-over-hand or inchworm-like, requires significant sub-diffraction spatial resolution (17, 19–22). Tracking experiments measure the velocity, run length (mean distance traveled before dissociation), dwell times between motions, and likelihood of stepping backwards. Comparing these data over a range of conditions, for instance varying the ATP concentration, with the predictions of kinetic models enables determination of the reaction pathway and limiting biochemical processes. Whether the experimental conditions can be broad enough to fully constrain the model and reach physiological conditions, such as millimolar ATP, depends on the spatial and temporal resolution of the experiment, and the speed of the motor. Dwell-times and velocities have straightforward saturable kinetics with ATP concentration, but the run length and probability of backwards stepping differ considerably among motors depending on the specifics of their kinetic schemes. Thus, theoretical modeling is an important tool for mechanistic interpretation of experimental data and hypothesis generation for behavior under different conditions, which leads to new experimental tests.

In order for the motor to exhibit directionality, either a mechanical or a biochemical asymmetry between the leading and trailing heads must be present (23). Biochemical asymmetry arises, for instance, in ATP binding and/or release of post-hydrolysis products, and it is believed to originate from inter-molecular strain between the two heads when they are both strongly bound to the filament. Due to their elastic connection, the two heads are under stress of opposite directions – the TH is pulled forward, and the LH is strained backward (24, 25). This can coordinate the ATPase cycles of the heads to cause efficient coupling between ATP utilization and stepping (24). A reduced rate of a biochemical step in the LH relative to the TH is thought to be the primary gating mechanism (26). Biochemical gating is not mandatory for kinetic schemes (27), but if it is absent, then some other directional bias must be present, for instance preferred detachment of the TH to initiate a step or promoted rebinding in the leading position, which can be confirmed by structural asymmetry of the catalytic cycle. Preferential rebinding in leading position of the stepping head can be considered a mechanical type of gating, but microscopic reversibility rules out a bias that remains in the absence of energy source (28). The mechanism of coordination adopted by a given motor is determined by structural features of the heads, intra-head linkers, and the mechanical load, and is tailored to the physiological role of the motor. Distinguishing between the distinct gating mechanisms by comparing predictions of kinetic models with experimental data, and relating them to structural details are therefore critical goals of research aimed at understanding intracellular transport.

These considerations have elicited a variety of theoretical approaches in order to obtain insights into the stepping mechanism (12, 29–40). One such method is to model motors as performing a random walk on a one-dimensional lattice. In this model, the motor is assumed to visit a number of biochemical and mechanical states in each position before completing a step. At each site *l* there are *N* intermediate chemical states, where *N* is usually determined either from biochemical kinetics, or by comparing theoretical predictions with experimental data. Formally, the probability of being in a state *i* at a location *l* along the track can be derived from the solution of an appropriate Markov chain or Master equation. Because cytoskeletal tracks are periodic, we can consider the directed motor motion as occurring on a periodic one-dimensional lattice (see Fig. 1).

**Figure 1:**
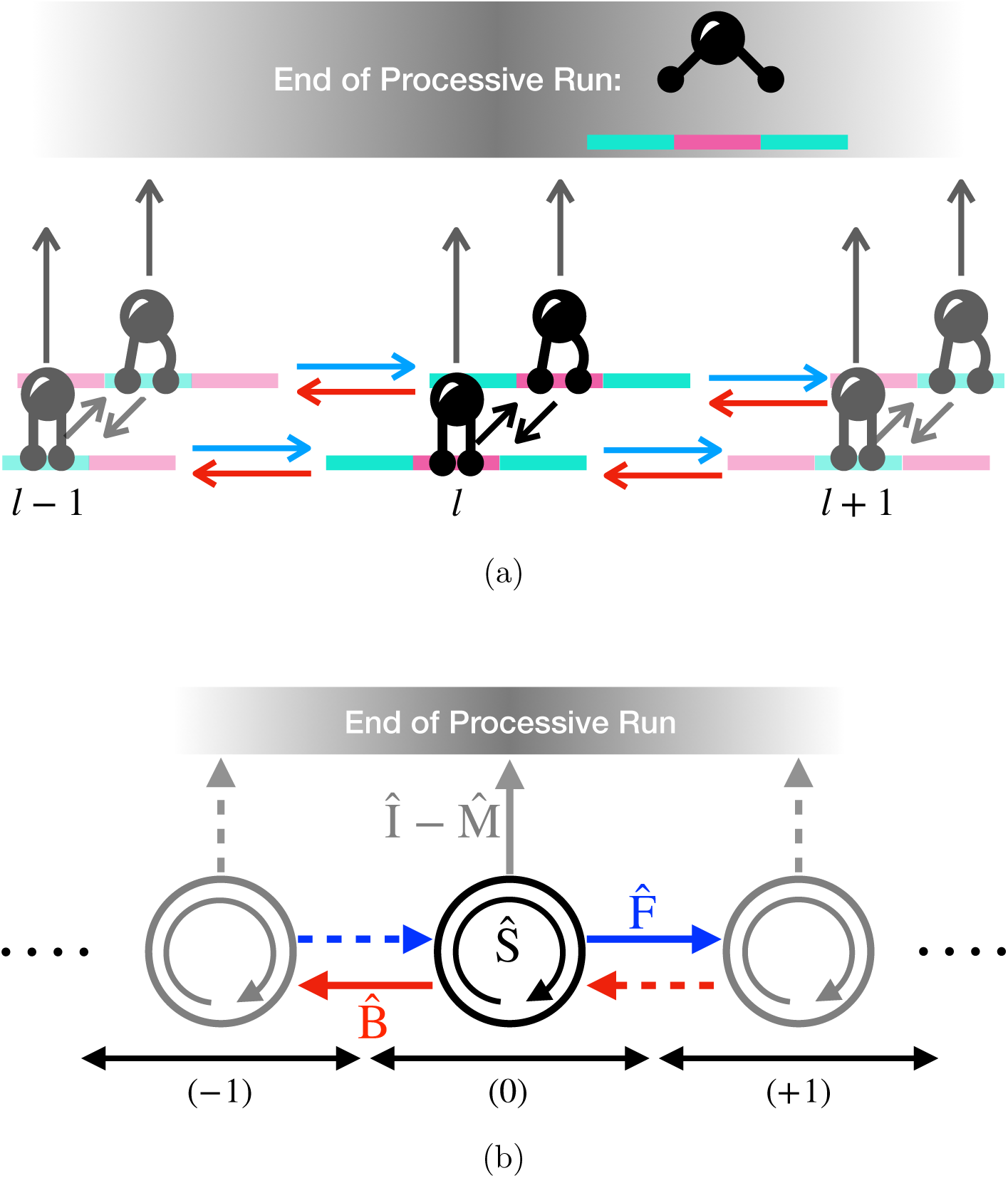
One dimensional network for a processive enzyme. (a) Two heads are attached to the same cargo (in black). The two heads are in shown in two conformations, with and without tension. The track is shown as green and magenta segments. Black arrows correspond to transitions between states at a fixed location along the filament (identified by the index *l*). Forward and backward steps are depicted as blue and red arrows, respectively. Grey arrows indicate the detachment from the track, which causes the end of a processive run. (b) Simplified representation of the more complete scheme, in which each symbol indicates a matrix associated with the corresponding transitions. The matrices 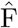 and 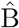 incorporate all the transitions associated with forward and backward steps, respectively. The matrix Ŝ contains all the rates related to changes in the conformation of the system at a fixed location along the track.

The solution of the model involves finding expressions that connect the microscopic rates describing biochemical and structural transitions with observable characteristics of motility, such as processivity, velocity, run length, and dwell-time (41–44). However, the kinetic scheme quickly becomes cumbersome, for instance with two heads that each explore 4 biochemical states (apo, ATP, ADP-P_i_ and ADP) bound-to and detached-from the track with asymmetry between the leading and trailing heads, 48 distinguishable intermediates are needed to describe the reaction scheme during a run, plus the detached state (an absorbing state) corresponding to run termination (45). Depending on the transitions allowed between these states, under the conservative assumption that only three transitions are possible per each head (involving either nucleotide binding/release, or attachment/detachment from the track) the number of kinetic rates is the order of a few hundreds (46). Of course, many of these rates are identical, or related to each other via thermodynamic considerations ensuring that the model does not result in a perpetual motion machine. The development of a formalism that is independent of the complexity of the network of connections between the accessible states, which is the aim of this work, facilitates a quantitative description and of the dynamical features of the model.

A way to simplify the description of this complex network is to define the *N* × *N* transition probability matrix, 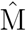, at any one location and to isolate the elements associated with transitions at a fixed site (Ŝ), from the forward steps (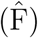), and backward steps (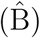). Periodicity imposes that the network of states, and the transition matrices be identical at every site along the track. Here we show that compact expressions emerge for the velocity, average and distribution of the run length (processivity), and probability of backward stepping for an arbitrary network of states. The expressions may be derived using matrix algebra, and can also be straightforwardly implemented for fitting experimental data. Our approach shares some similarities with the strategies adopted in (27) and (47) to extract the velocity and processivity for kinetic models of myosin VI, and X, respectively. However, our results have the benefit of an analytical solution which holds regardless of the underlying structure of the kinetic network.

In the next sections, we first discuss the main results of the paper, and illustrate them in the context of a simple one-dimensional random walk. We then consider a modification of the model, which allows it to account for steps of different sizes. Finally, we illustrate the utility of the method with a case study of the processivity of dimeric myosin VI.The literature contains conflicting conclusions about biochemical and mechanical gating in myosin VI (25, 27, 48, 49), and we show that our the theoretical solution to the model can be used to reveal the gating mechanism, the pathway for backward stepping, and the energy consumption of the motor.

For clarity of presentation, we relegate the formal derivation of the analytical results and the details of Kinetic Monte Carlo (KMC) simulations to the Supplementary Information (SI).

## Results

### Processivity and Probability of Forward Stepping

In order to present the major results of the paper, some new nomenclature is required. First, we distinguish *transitions* from *steps*. A transition occurs whenever the biochemical state or the structure of the motor changes. A step is a transition associated with a forward or backward displacement along the track. We use the index *x* to identify transitions, *n* for the total number of steps (backward and forward), *f* and *b* refer to the counts of the forward and backward steps, respectively. A “pathway” is a series of transitions, whereas a “trajectory” refers to a collection of steps. In the discussion of the main results and in the SI we introduce a number of different symbols defined in Table S1.

Let *N* be the number of distinguishable biochemical states accessible to the motor. The *N* states are assumed to be bound to the track, which means that at least one member of the dimeric motor complex is attached to the filament. If both of the heads simultaneously detach from the track, the motor dissociates from the filament; we consider this state to be “absorbing,” implying that there are no further transitions from it and the run cannot continue.

The *N*-dimensional vector 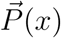 gives the probabilities of occupying any of the *N* states after *x* transitions, regardless of the location along the track. In a Markov chain, the probability vector 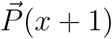 is obtained multiplying 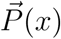 by a transition probability matrix, whose *N* ^2^ entries define the probabilities of changing location (stepping) and/or the biochemical state of the motor. We consider four matrices: Ŝ contains all the transitions that occur at a fixed location on the track, 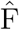 and 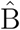 include all of the forward and backward steps, respectively, and 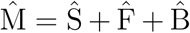 accounts for all possible transitions between bound states. Because of the periodicity of the track, 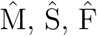, and 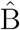 are identical at any site along the filament, therefore, for an infinitely long track, 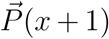 is given by (see SI for details),

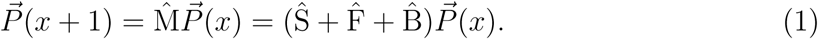

The probability of being bound to the track is given by the sum of the probabilities of being in any of the *N* biochemical states, that is 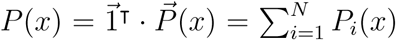, where 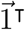 is an *N*-dimensional vector of ones. At 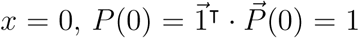, but as the state of the system evolves the motors dissociate from the filament, therefore *P* (*x*) ≤ 1, and *P* (*x*) → 0 for *x* → ∞.

We are interested in obtaining the probability ℙ (*n*) that the motor takes exactly *n* steps (forward or backward) before detaching from the track. We show in the SI that,

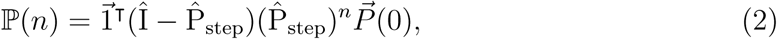

where,

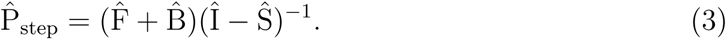

The matrix 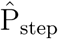 accounts for all the possible pathways ending with a forward or backward step [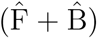 in Eq. 3] after an arbitrary number of transitions has occurred at a fixed location [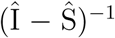 in Eq. 3, see SI for details]. The expression in Eq. 2 has a simple physical meaning. It is clear that (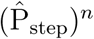 includes all the trajectories in which exactly *n* steps take place, regardless of the number of actual transitions. The matrix 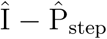 is the probability of detachment from the track before the occurrence of any other step, but after an arbitrary number of transitions has taken place. The product of these terms [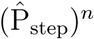 and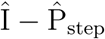] is ℙ (*n*) given in Eq. 2.

It may be easier to understand Eq. 2 by considering the case for *N* = 1 depicted in Fig. 2. When *N* = 1, the transition probability matrices 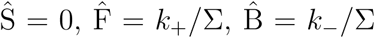, and 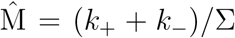 are all scalars, with Σ = *k*_+_ + *k*_*-*_ + *γ* being the sum of all the rates. It is easy to show that 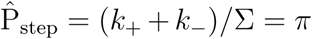, and that 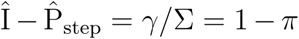, where *π* is the probability of stepping either forward or backward, and 1 *- π* is the probability of dissociating from the track. Plugging these expressions in Eq. 2 (in the *N* = 1 case 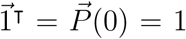) we find that ℙ (*n*) = (1 *- π*)*π*^*n*^, a well-known and intuitive result. Therefore, Eq. 2 generalizes the *N* = 1 case to an arbitrary number of biochemical states.

**Figure 2:**
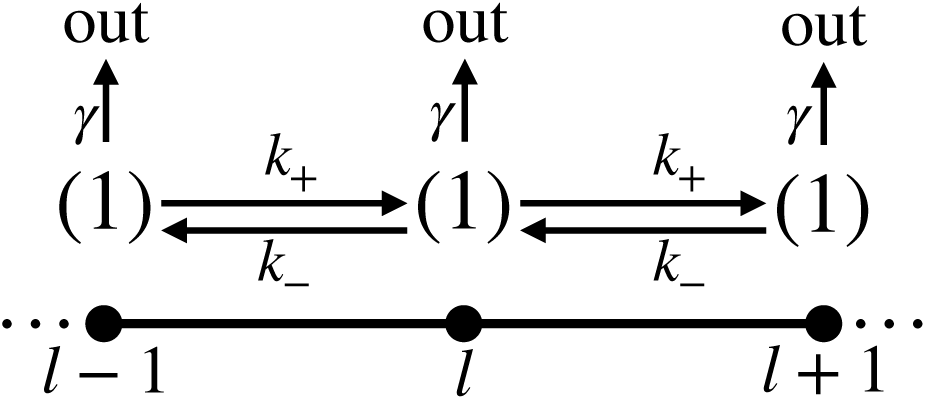
Molecular motor without intermediate states. Detachment, forward stepping, and backward stepping and stepping occur only from state (1), with rates *γ, k*_+_, and *k*_*-*_, respectively.

Once the distribution ℙ (*n*) is known, it is straightforward to derive the average number of steps completed before detachment. After some algebraic manipulations (see the details in the SI), the result is,

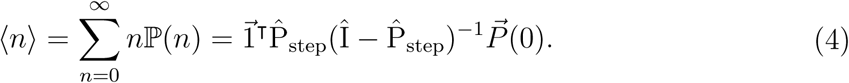

Again, it is useful to consider the *N* = 1 case, in which Eq. 4 becomes ⟨*n*⟩ = *π/*(1 *- π*), another well-known result.

Analogously, we derived the probability of detaching after *f* forward steps have been completed, regardless of the number of backward steps or transitions at a fixed site. As shown in the SI, we obtained the average number of forward steps before detachment as,

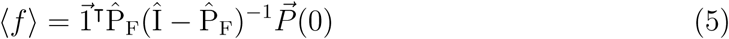

With

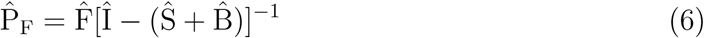

The physical reasoning is the same as before. The matrix 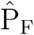 accounts for all pathways ending with a forward step 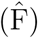 after an arbitrary number of backward steps and transitions at a fixed location have occurred 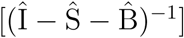. Similarly, the average number of backward steps completed before detachment, regardless of the number of forward steps or total transitions, is,

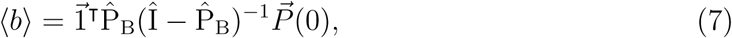

with

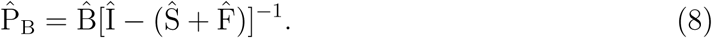

Note that Eq. 7 is the same as Eq. 5 after swapping 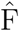 and 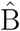.

As before, it helps to think about the *N* = 1 case, in which 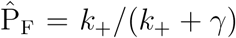 and

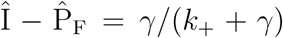, leading to ⟨*f* ⟩ = *k*_+_*/γ*. Similarly, we find that when *N* = 1, ⟨*b*⟩ = *k*_*-*_*/γ*.

We define the probability of stepping forward as the average number of forward steps divided the total number of steps completed before detachment,

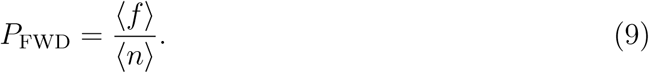

As we show in the SI, ⟨*n*⟩= ⟨*f* ⟩ + ⟨*b*⟩.

Finally, the average run length is given by,

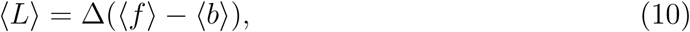

where Δ is the average step-size of the motor. When *N* = 1, *P*_FWD_ = *k*_+_*/*(*k*_*-*_ + *k*_+_), and ⟨*L*⟩ = Δ(*k*_+_ *- k*_*-*_)*/γ*, which are the expected results.

### Average Run Time

Let ⟨*τ* ⟩ be the average run time of the motor, that is the average duration of a processive run. Here, we briefly outline how to obtain an expression for ⟨*τ* ⟩ independent of the number of biochemical states and the connectivity between them. We begin defining the vector 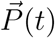, which incorporates the probability of being at any of the *N* states bound to the track at time *t*, anywhere along the filament. The evolution in time of 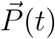 is obtained by solving the Master equation,

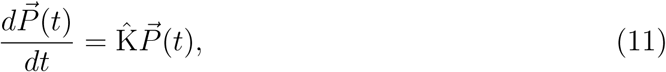

where 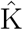 is the *N* ×*N* rate matrix. We ought to point out that 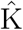 includes the transitions towards the absorbing state. Using a classic result (50), the run time (or mean first passage time to detachment) is,

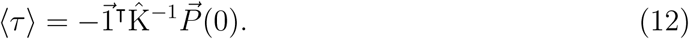

### Velocity

In order to obtain an expression for the average velocity, we introduce the probability *p*(*L, τ*) *dτ* of detaching after a run length *L* and a run-time *τ*. The most direct definition of the average velocity is *v* = ∑_*L*_ ∫*dτ L/τ p*(*L, τ*) = ⟨*L/τ* ⟩. Such an approach has been taken for a simple analytical model for kinesin; an analytical expression for *p*(*L, τ*) can be found in this case (38). Alternative definitions of *v* are (i) the limit lim_*t→∞*_ *d*⟨*l*(*t*) ⟩*/dt* (29, *41)*, where *l*(*t*) is the location of the motor at time *t*, and the average is performed over the ensemble of motors that are still bound to the track at time *t* (51),(ii) the “instantaneous” velocity, taken as the average step-size multiplied by the rate of stepping, and (iii) the ratio between the average run length and the mean run time ⟨*τ* ⟩,

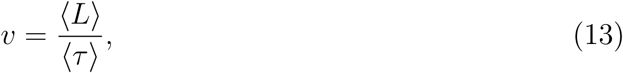

which is the definition adopted here. We note that both ⟨*L/τ* ⟩ and ⟨*L*⟩*/* ⟨*τ* ⟩ can be obtained from experiments. In general, their values do not coincide, though we find empirically that they become closer as the processivity of the motor increases (see Fig. S9d).

### Multiple Stepping Mechanism

So far, we have assumed that all the different stepping pathways displayed by a motor are characterized by an identical average step-size (Δ in Eq. 10 is the periodicity of the track). However, this is not always the case. For instance, dynein and myosin VI display broad step-size distributions (22, 52). In particular, myosin VI (discussed in detail later) combines hand-over-hand with inchworm-like displacements (52). The theory presented here can be generalized in order to incorporate step-size variability. Suppose that a dimeric motor exists in two states, D (as in “distal”), with the two heads separated by a repeat of the filament, and C (as in “close”), in which the motors are bound to adjacent sites. This scenario is represented in Fig. 3, and is inspired by a model for myosin VI proposed by Yanagida and coworkers (52–54) (discussed in further detail in the next section). The number of D and C states are *N*_D_ and *N*_C_, respectively, and *N* = *N*_D_ + *N*_C_ is the total number of bound states available to the motor. Let 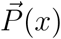 be the *N*-dimensional vector describing the probabilities of occupying the *N* different states anywhere along the filament. The matrices ŜD and ŜC contain the transitions within states D and C, respectively, explored without changing location along the track. The forward and backward transitions associated with hand-over-hand steps are described by the matrices 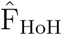 and 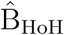, respectively, and connect states D at different filament sites. Inchworm steps are accounted for by the matrices 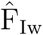 and 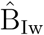, and connect D and C states. Let Ŝ = ŜD + ŜC, we show in the SI that, under the assumption of periodic and infinite track, the following equation holds,.

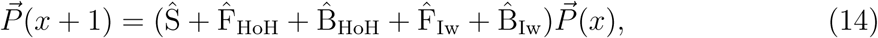

which becomes identical to Eq. 1 upon defining 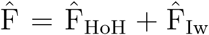 and 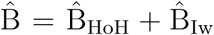. Similarly to what we have done before, we can compute the average number of forward and backward hand-over-hand and inchworm steps, ⟨*f*_HoH_⟩, ⟨*b*_HoH_⟩, ⟨*f*_Iw_⟩, and ⟨*b*_Iw_⟩, with ⟨*n*⟩ equal to the total number of steps. It follows that we can define a probability of taking a forward hand-over-hand or inchworm step,

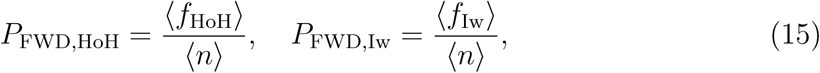

and the average run length is,

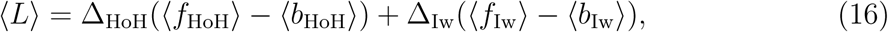

where the inchworm step-size is Δ_Iw_ = Δ_HoH_*/*2.

**Figure 3:**
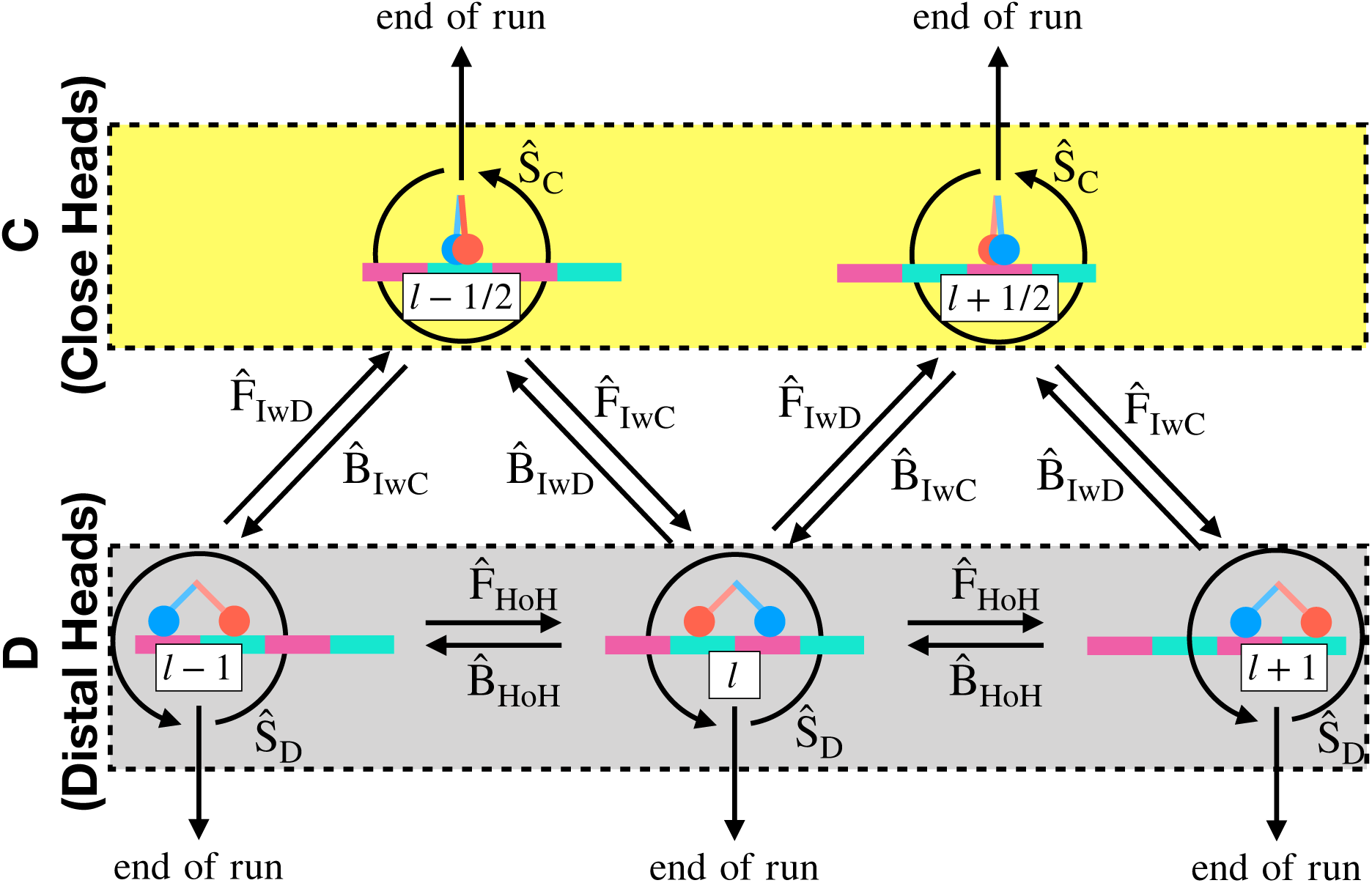
Combination of hand-over-hand and inchworm-like steps. The two heads forming the dimer are shown in red and blue, the track is in green and magenta. The bottom three states are of type D, with the two motors occupying sites separated by a filament repeat. In the case of states C (upper two states) the two heads are closer to each other. Hand-over-hand steps connect D states at different sites along the filament, and the corresponding transition probabilities are contained in the matrices 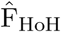 and 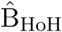. There are two types of inchworm-like steps, both of which connect states D with C. The motor could either step from D to C (matrices 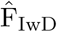 and 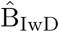) or from C to D (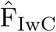 and 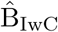). Transitions that do not change the location of the motor are described by the probability matrices ŜD and ŜC for states D and C, respectively. Dissociation from the filament may occur from either D or C.

### Observables

From the kinetic mechanism, we can compute the average number of times that any specific event occurs. For instance, we may be interested in monitoring the energy consumption of the motor, which can be calculated by counting the average number of ATP molecules hydrolyzed per processive run. Let *q* be the count of the event of interest. In order to compute ⟨*q*⟩, we collect in the matrix 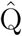 all the transitions corresponding to the occurrence of the event of interest, and 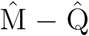 is the probability matrix associated with all the remaining transitions. We show in the SI that,

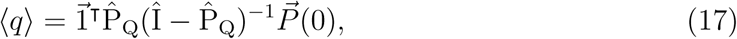

with,

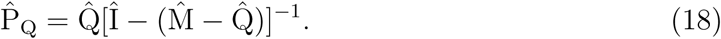

There is a similarity between this expression and the average number of steps previously discussed. For instance, let 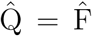, in this case 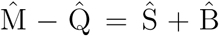 per definition, and 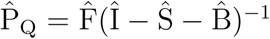, which is the same as 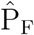 in Eq. 6. Therefore, when 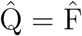 Eq. 17 is the same as Eq. 5. Similar arguments show that from Eq. 17 we recover Eq. 4 if 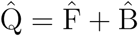, and Eq. 7 if 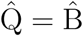.

## Myosin VI Gating Mechanism

### Myosin Stepping Pathways

Myosins are molecular motors typically composed of three structural regions with distinct functions: (i) a head domain, which hydrolyzes ATP and binds actin, (ii) the oblong lever arm, which amplifies the small structural transitions occurring in the myosin head, and (iii) a tail domain responsible for multimerization and cargo binding.

Processive myosins (M) are tightly bound to the actin filament (A) when they are in complex with ADP (AM*·*D) or in the apo state (AM) (13–15). ATP binding to the motor induces detachment from actin. After ATP hydrolysis myosin binds actin tightly, undergoes the power stroke (a structural transition that propels the lever arm forward), and releases the P_i_. The re-priming stroke occurs upon ATP hydrolysis, when the motor is dissociated from the actin filament, and rotates the lever arm to the pre-stroke position.

Dimeric myosins walk predominantly by a hand-over-hand mechanism (9, 17, 19, 20), in which the trailing head (TH) detaches from the filament, overtakes the bound head, binds to a new actin subunit, and becomes the new leading head (LH). As discussed in the Introduction, this stepping mechanism is facilitated by two types of processes, (i) biochemical gating, and (ii) mechanical gating. The former ensues from the inter-head tension between the two bound motors, and is expected to facilitate the detachment of the TH before the LH by modifying the rates of nucleotide binding and release of the two heads. The second is structural, due to the combined effect of the power stroke in the leading head and re-priming stroke of the detached head, which favor the reattachment of the stepping head in the leading position.

However, exceptions include, (i) backward steps, in which the LH detaches first and moves rearward (52), and (ii) inchworm-like steps (52), leading to transitory states in which the two motors bind at adjacent actin sites. Also, (iii) the inter-head tension experienced by the two actin-bound motors may affect the stability of the acto-myosin complex both in apo and in ADP-bound state (25), hence it may facilitate the *spontaneous* detachment from the filament (i.e. occurring prior to ATP binding). Finally, (iv) a stepping head may “stomp,” and rebind to the same location on actin from which it detached (55, 56). Stomps may correspond to futile cycles, in which an ATP molecule is hydrolyzed without motor movement.

Ideally, a model for myosin should incorporate all of the above features, and their measured occurrence should emerge as a natural consequence of the model. The method presented in this manuscript provides a straightforward route to a comprehensive model which encompasses all of these stepping modes.

### Application to Myosin VI

We apply the theory to myosin VI, the only member of the myosin superfamily that is known to move towards the pointed end of the polarized actin filament (57), and a processive motor that is also known to take frequent backward and inchworm-like steps (52). Despite a number of insightful studies, the nature of the gating mechanism in some of the myosin motors, including myosin VI, continues to be unresolved. It has been suggested that ADP release is slower in the LH than in the TH (ADP-release gating) (27, 49). Alternatively, it was proposed that ATP binding occurs more slowly in the LH (ATP-binding gating) (25, 48, 58). In addition, recent experiments have suggested that the probability of forward stepping (compared to backward stepping) for myosin VI depends on the concentration of ATP (54), a property anticipated for either ADP release, or ATP binding gating. Thus, our model, which naturally includes all of the stepping modes provides clarity to the debate by accounting fully for the published experimental data (27, 54). All the parameters and rates of the model are described in Table S2.

The model for myosin VI dimers is sketched in Fig. 3, where we show both states of type C (as in “close”) in which the heads are bound to adjacent actin sites, and states of type D (as in “distal”), with the two motor heads separated by a filament repeat (≈ 36nm). This type of model has been proposed by Yanagida and coworkers in a number of remarkable experimental studies of myosin VI processivity (52–54).

Figure 4 shows the biochemical states explored by each individual myosin head. We ignore the collision complexes – all the nucleotide binding events are considered to be pseudo-first-order – and we assume that ATP binding induces a fast detachment from actin. Furthermore, the transition A+M*·*T → AM*·*D combines (i) ATP hydrolysis, (ii) formation of the transitory AM*·*D*·*P_i_ complex, and (iii) actin-activated phosphate release. We ignore the slow phosphate release from the detached motor in the one-head-bound (1HB) state.

**Figure 4:**
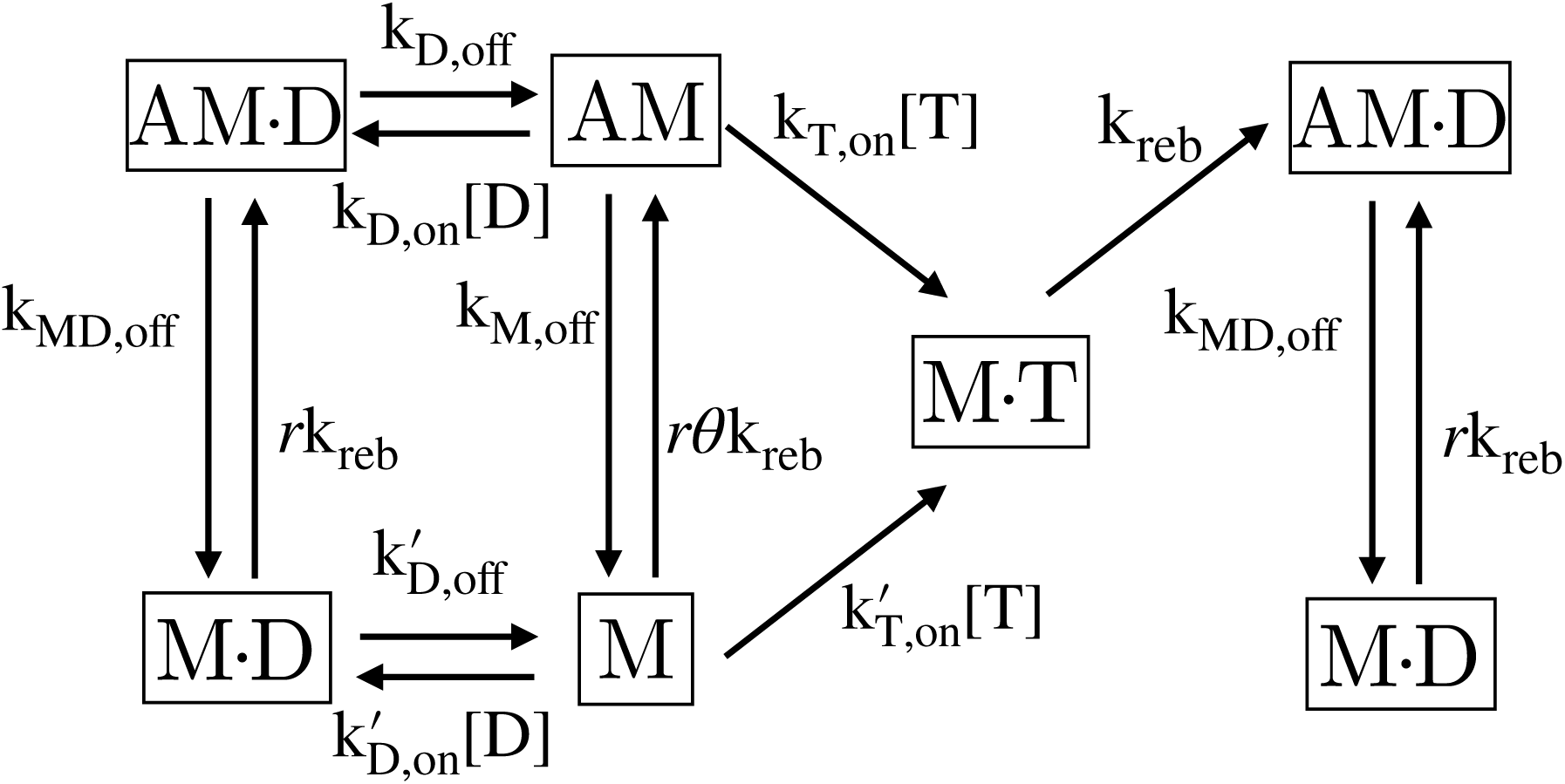
States accessible to a single myosin. The scheme is a simplified representation of the myosin cycle shown in (13–15). A is actin, myosin is M, ADP and ATP are shown as D and T, respectively, and P_i_ is orthophosphate. All the rates are described in Table S2. ATP binding to actomyosin is assumed to lead to a fast detachment of the motor from the filament. Rebinding is a combination of ATP hydrolysis, binding to actin, and releasing the phosphate. ATP binding is assumed to be irreversible. *r* is a parameter describing the enhanced rate of actin binding in the ADP-bound and apo state compared to the ATP-bound state. *θ* relates the rate of actin binding in the apo and ADP-bound state.

For the dimeric motor, *N*_D_ = 16, with 4 states in which both heads are actin-bound (two-head-bound, or 2HB, states), and 12 in which only one head is attached to the track (1HB states). In the C states, experiments suggest that the two heads step with equal probability (53), and as a consequence there are only 3 states with 2HB, and 6 states with 1HB, for a total of *N*_C_ = 9. The total number of actin-bound states is *N* = 25 (59).

We assume that there is no tension between the heads when the motors are sitting at adjacent sites, corresponding to the C state. Thus, biochemical gating is not actin in C, but tunes the relative rate of ADP release from and ATP binding to the LH of a 2HB state D: 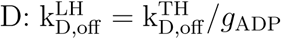 and 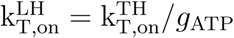, where *g*_ADP_ and *g*_ATP_ are the factors identifying the strength of ADP-release and ATP-binding gating. When the heads are bound to adjacent subunits (states C) we adopt the rates 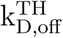 and 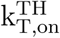 for both heads.

The rate for the A+M.T→A.M.D transition is k_reb_ = 57s^*-*1^ (27). The binding rate of a dissociated ADP-bound myosin head is *r*k_reb_, where the factor *r* is an adjustable parameter. For an apo head, we use *rθ*k_reb_, with *θ* set via thermodynamic considerations (see later and SI). We expect *r* > 1 because of the high duty-ratio of myosin VI (14).

The motor can stomp, complete a hand-over-hand step (from state D to D) of size Δ_HoH_ = 34 nm (27, 49), or a inchworm-like step (D→C or C→D) with Δ_Iw_ = 17 nm. The detached head in a 1HB state may bind to three possible sites along actin: in leading position, trailing position, or adjacent to the bound head. The relative likelihood of these three 2HB geometries depends on the orientation of the lever arm of the two heads when the free motor binds actin. Let *g*_MEC_ be the strength of the mechanical gating due to either the re-priming stroke in the detached head or the power stroke of the attached head. The probability of attaining a 2HB state when both heads are in their preferred orientation is enhanced by a factor 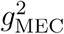. If only one of the lever arms must rotate away from the preferred angle the factor is *g*_MEC_, whereas there is no enhancement factor if the dimer geometry at the time of binding actin is frustrated in both the heads simultaneously. The lever arm of an actin-bound head is always in the post-power-stroke orientation, that is it points forward. In contrast, for the lever arm of the free head, we distinguish two scenarios, depending on whether the detached head of a 1HB state is or is not bound to ATP. The scenarios are portrayed in Fig. S2 of the SI.

(1) If during a step ATP is hydrolyzed, the free head is in primed state, then the lever arm points backwards on actin binding, that is it preferentially binds actin in leading position (see Fig. S2a). Therefore, we adopt the following sets of rates for binding actin as the leading head (*l*_MT_), in trailing position (*t*_MT_), or in the vicinity of the bound head (*c*_MT_, in state C),

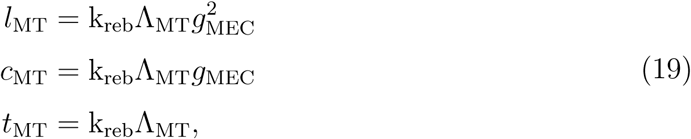

where 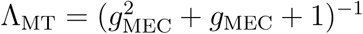.

(2) If the detached motor is in the apo or ADP-bound state, the lever arms of both heads (the one bound to the track and the free one) are in the post-stroke position, and thus the C state allows both heads to maintain their favored lever arm orientation upon rebinding, resulting in a landing rate proportional to *r*k_reb_*g*_MEC_*/*(*g*_MEC_ +2) (see Fig. S2b). A nucleotide-free or ADP-bound dissociated head binds actin in state D (either as the LH or as the TH) with rate proportional to *r*k_reb_*/*(*g*_MEC_ + 2), because one of the two heads must rotate the lever arm away from the preferred orientation in order to accommodate the 2HB geometry (see Fig. S2c).

The rates for spontaneous detachment and the rates for ADP-bound and apo state binding to actin are related to each other via thermodynamic considerations (see details in the SI, Fig. S3-S4). We find that the following conditions must hold.

(1) The ADP dissociation constant for an actin-bound head (*K*_D_) is the same regardless of whether the motor in 1HB or 2HB state, in D or C state, in leading or trailing position (see Eq. S24).

(2) The ADP dissociation constant for a myosin head detached from actin 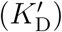 is related to *K*_D_ through a function of the rates for spontaneous detachment and acting binding in the apo or ADP-bound states (see Eq. S24).

(3) Finally, spontaneous stepping cannot on average lead to directional movement (see Eq. S25).

In accordance with these constraints, we adopt the following relationships between the rates of spontaneous detachment and the rates for binding actin in the ADP-bound or apo state,

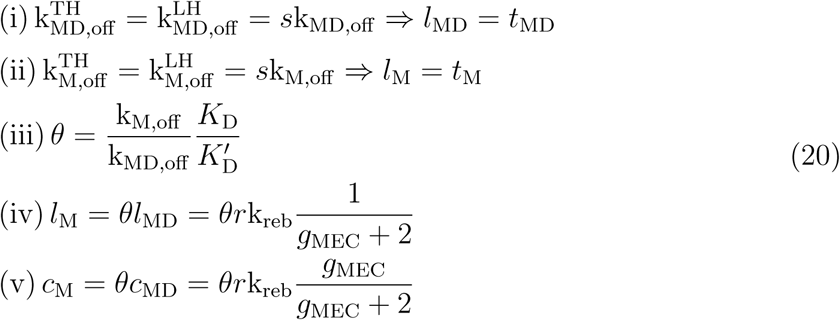

In relationships (i) and (ii) we assumed that the rates for spontaneous detachment from the LH and TH are the same, and that they are equal to the corresponding rates for a single head (*k*_MD,off_ and *k*_M,off_) times a fitting parameter *s*, which is assumed to be the same for both the apo and ADP-bound state. Because tension is known to facilitate the detachment of the myosin heads (25), we expect *s* > 1. Spontaneous stepping cannot lead to directional movement, therefore 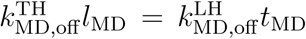, and similarly for the apo state (see Eq. S25). The identities in relationships (i) and (ii) after the “⇒” symbol follow. For a motor in the C state or for the bound head in the 1HB state there is no tension between the heads, therefore we use the rates k_M,off_ and k_MD,off_ for spontaneous detachment. The third identity [(iii) in Eq. 20] defines the parameter *θ*. Because of the choices made in identities (i) and (ii), *θ* is obtained as a combination of quantities that are directly measured in experiments. The fourth and fifth identities [(iv) and (v) in Eq. 20] relate the rates for binding actin in the apo and ADP-bound states via the parameter *θ*, as required by the constraints on the model (see SI).

In order to complete the model, we need the initial condition, 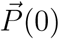 We assume that the two detached myosin VI heads behave independently, and that they exist in three possible states: apo, ATP-bound (which we assume equivalent to a ADP*·*P_i_), and ADP-bound. Movement along actin begins when both heads bind to the filament. The equations used to establish the initial condition are given in the SI.

We computed the (i) run length, (ii) velocity, and probability of taking forward (iii) hand-over-hand and (iv) inchworm steps as a function of ATP concentration using the approach described above. Our predictions are compared to the run length and velocity data from Elting *et al.* (27) (Figs. 5a-5b), and the probability of taking hand-over-hand and inchworm steps is compared to the probability for long and short forward steps measured by Ikezaki *et al.* (54) (Figs. 5c-5d). The parameters adopted are reported in Table 1. We adjusted four quantities: *(i) g*_ADP_, (ii) *g*_MEC_, (iii) *s*, and (iv) *r*. We find that letting *g*_ATP_ free to vary does not significantly change the quality of the fit, and *g*_ADP_ remains the dominant biochemical gating mechanism (see caption of Table 1). The model recovers quantitatively the results from the two different groups (Fig. 5). The main difference is the average run length, which can at least in part be explained because we neglected the finite size of the actin filament (27).

**Table 1:** Parameters used in the myosin VI model. The left column shows the name of the parameter or the corresponding transition. The parameters in the central column refer to transitions of the TH or LH in a 2HB state, transitions involving the bound and unbound myosin in the 1HB states, or parameters used to determine the initial state (zero head bound, or 0HB). The last column gives the numerical value of the parameter. The parameters marked with a * are taken from (27), and the † symbol refers to the mutant T406E of (14). *θ* was obtained using the listed ADP-release rates and spontaneous detachment rates, and the ADP-binding rates from (14). The values in bold are fitted to the data (see Methods section). After the minimization we obtained *χ*^2^*/n*_data_ ≈ 11.83, where *n*_data_ is the number of data points considered. Allowing *g*_ATP_ to vary does not significantly alter the result of the fit [*g*_ATP_ ≈ 2.99, with *g*_ADP_ ≈ 8.54 and *χ*^2^*/n*_data_ ≈ 11.73]. As described in the main text, 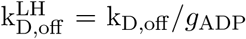 and 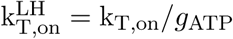. In addition, 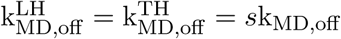, and 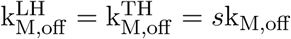.

| scheme | rate | applied to | value |
| --- | --- | --- | --- |
| AMD→AM+D | $k_{D,\text{off}}$ | 2HB(TH) or 1HB | $5.1/s^\star$ |
| AMT→A+MT | $k_{T,\text{on}}$ | 2HB(TH) or 1HB | $0.016/(s \cdot \mu M)^\star$ |
| AMD→A+MD | $k_{MD,\text{off}}$ | 1HB | $0.06/s^\dagger$ |
| AM→A+M | $k_{M,\text{off}}$ | 1HB | $0.005/s^\dagger$ |
| A+MT→AMD | $k_{\text{reb}}$ | 1HB | $57/s^\star$ |
| MD→M+D | $k'_{D,\text{off}}$ | 1HB or 0HB | $6.4/s^\dagger$ |
| M+T→MT | $k'_{T,\text{on}}$ | 1HB or 0HB | $0.27/(s \cdot \mu M)^\dagger$ |
| MDP<sub>i</sub>→MD+P<sub>i</sub> | $k'_{P_i,\text{off}}$ | 0HB | $0.04/s^\dagger$ |
| parameter |  | applied to | value |
| $g_{\text{ADP}}$ | | 2HB(LH) | **6.01** |
| $g_{\text{ATP}}$ | | 2HB(LH) | 1.0 |
| $g_{\text{MEC}}$ | | 1HB or 0HB | **15.6** |
| $s$ | | 2HB | **9.93** |
| $r$ | | 1HB or 0HB | **1.62** |
| $\theta$ | | 1HB | 0.117 |

**Figure 5:**
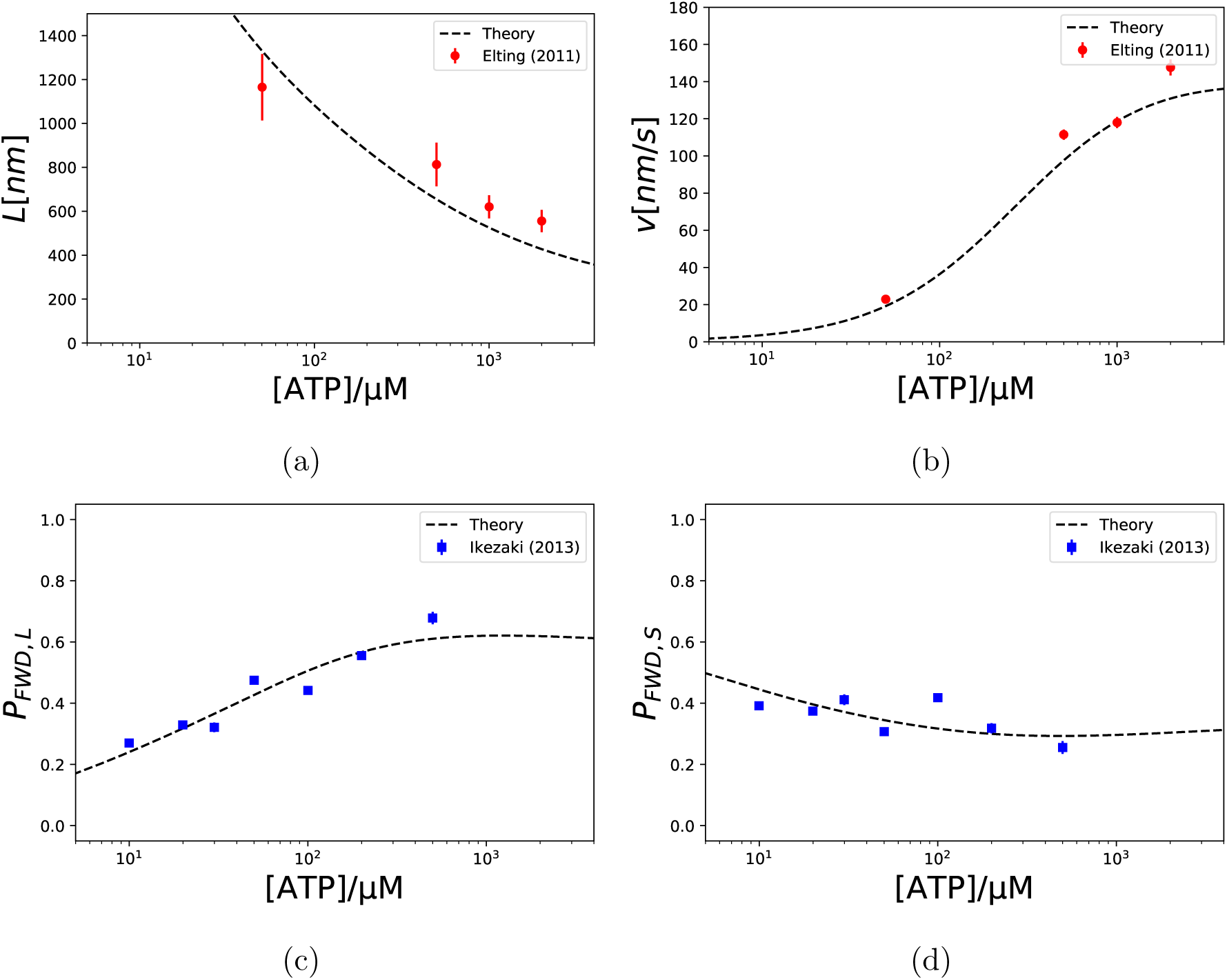
Comparison of theoretical results to experiments on myosin VI. The red points and blue squares were digitized using *WebPlotDigitizer* (61) from Fig. S6c-d of Elting *et al.* (27) and Fig. 3B of Ikezaki *et al.* (54), and the dashed lines come from the theory, after fitting the parameters (see Table 1). (a) Run length. (b) Velocity. (c) Probability of taking hand-over-hand and (d) inchworm-like forward steps.

### Goodness of Fit

We also investigate whether the fitted parameters are well constrained. Figure 6 shows the relative changes of the squared residual error obtained when each pair of parameters is modified. ATP and ADP gating have a weak correlation, but an optimal fit requires that biochemical gating is predominantly due to more rapid release of ADP from the TH (Fig. 6a). We note that *rθ* < 1, which indicates that binding actin in the apo state is slower than in the post-ATP-hydrolysis conformation. While this is possible, the upper bound for the parameter *r* is not tightly constrained, so it may be possible to find a combination of parameters for which *rθ* > 1. We suggest that data at lower ATP concentrations (in particular for the run length) could help constrain the parameters.

**Figure 6:**
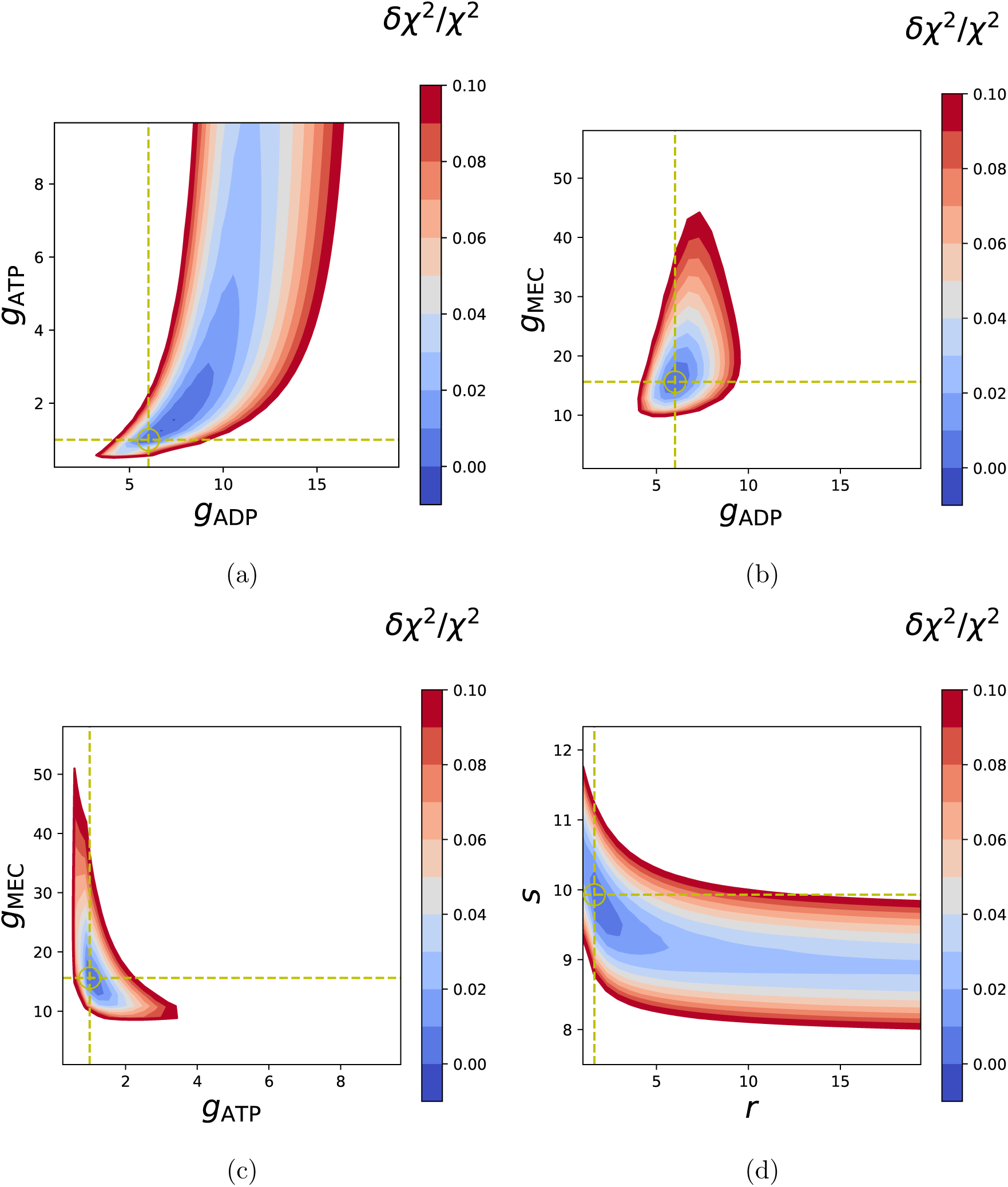
Dependence on the parameters of the quality of the fit of the theory to experiments on myosin VI. The four panels show different pairs of fitted parameters. The color code reports the relative changes of *χ*^2^ (defined in the Methods section) obtained when two parameters are changed – the remaining ones are kept at the value determined with the fit. The white area indicates a region in parameter space where the relative change of *χ*^2^ is larger than 10%. The yellow dashed lines and circle identify the minimum found with the fit. (a) ADP versus ATP gating. (b) ADP versus mechanical gating. (c) ATP versus mechanical gating. (d) *r* versus *s*, where *r* is the enhancement of the rate for binding actin in the ADP-bound and apo state compared to the post-hydrolysis conformation, and *s* is the enhancement of the rate for spontaneous detachment of the two heads when they are subject to inter-head tension in the 2HB, D state.

### Predictions

In Fig. 7, we show the theoretical predictions. First, the majority of backward steps are inchworm-like (Fig. 7a, red, dashed line), which is in agreement with myosin VI step-size distribution (52), as well as with earlier measurements reporting a load-independent average backward step-size of ≈ 11 nm (60). (Note that the measured step-size may depend on the geometry of the motor and the location of the probe.) Second, the majority of backward steps are spontaneous (Fig. 7b, red, dashed line), which agrees with the analysis performed by Ikezaki *et al.* (54). As expected, the forward steps are predominantly fueled by ATP (Fig. 7b, solid, blue line). Stomps are spontaneous at low ATP concentrations (Fig. 7b, dashed, grey line), but for [ATP] > 2 *µ*M these futile cycles waste a molecule of ATP (Fig. 7b, solid, grey line). Third, we investigate whether myosin VI uses ATP parsimoniously. We make the following considerations. Backward steps effectively waste ATP, because after a rearward movement it takes an ATP-driven forward step to restore myosin to its initial position. Thus, ATP-fueled backward steps (⟨*b*⟩_ATP_) dissipate the energy from 2 ATP molecules per backward step, and the imbalance between spontaneous backward and forward steps (⟨*b - f* ⟩_spo_) accounts for the net expenditure of 1 ATP per net spontaneous backward motion to restore the position. In addition, ATP is dissipated also when the motor stomps after hydrolysis (⟨*s*⟩_ATP_). These considerations allow us to define the following function, *W*, which monitors the proportion of futile ATPase cycles,

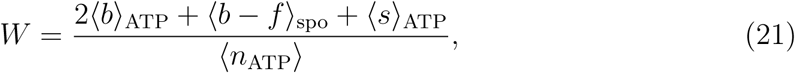

where ⟨*n*_ATP_⟩ is the total number of ATP molecules hydrolyzed during the processive run. In Fig. 7c the function *W* has a peak around 10 *µ*M, where both the imbalance of forward and backward spontaneous steps and ATP-driven stomps combined to waste approximately half of the ATP hydrolyzed. At high ATP concentrations (≈ 1 mM), ≈ 13% of the hydrolysis cycles are unproductive, predominantly because of ATP-induced stomps (Fig. 7c, grey curve). This result constitutes a novel prediction, underscores the importance of including stomps to estimate the energetics of motors, and sets an upper bound on the proportion of productive ATP hydrolysis cycles.

**Figure 7:**
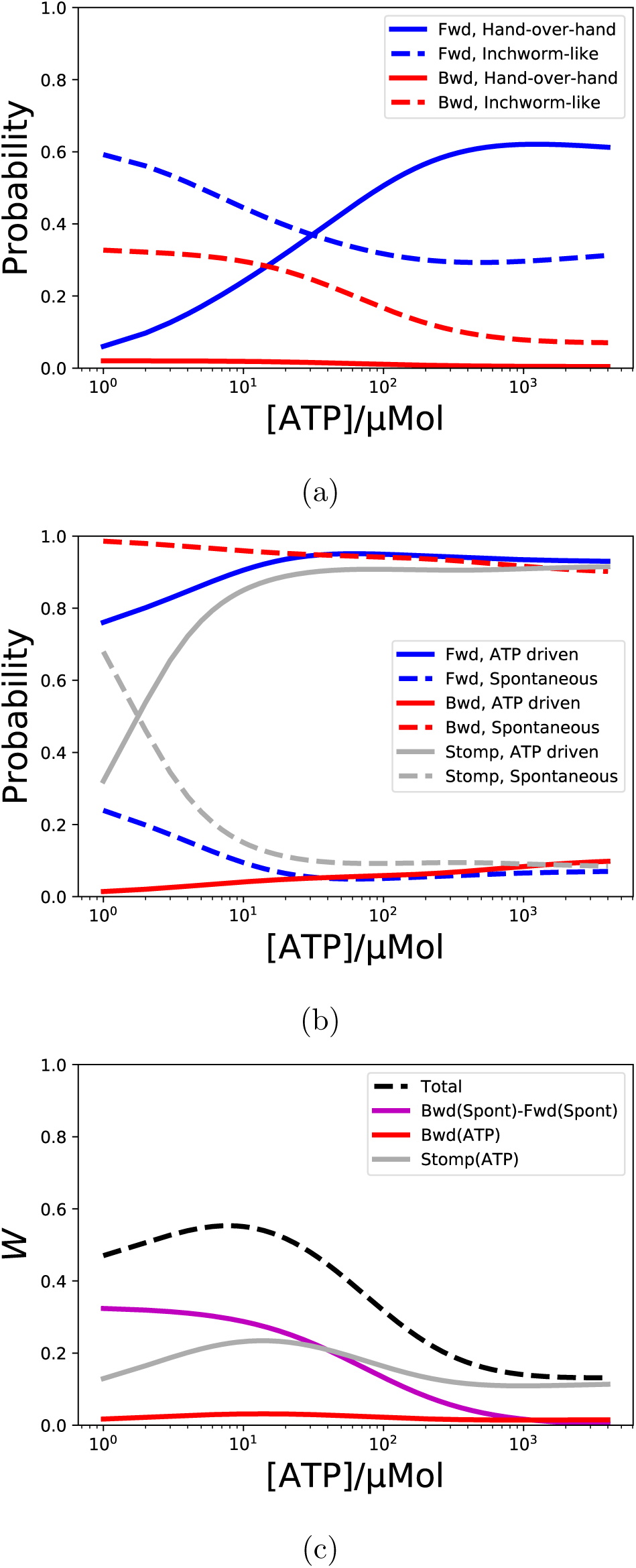
Theoretical predictions. In all figures, blue lines refer to forward steps, backward steps are in red, grey lines are stomps. (a) forward and backward stepping pathways. The continuous lines indicate hand-over-hand steps, inchworm-like steps are shown as dashed lines. (b) Conditional probabilities of taking a forward step, backward step, or stomp fueled by ATP (continuous lines) vs spontaneous (dashed lines) given that a forward step, backward step, or foot stomp occurs, respectively. (c) Waste probability *W* (see Eq. 21) as a function of [ATP]. The black, dashed line indicates the value of *W*, which is the cumulative result of imbalance of spontaneous steps (magenta), and ATP-consuming backward steps (red) and stomps (grey).

## Conclusions

We presented a theory to predict the number of steps taken by a processive motor on a periodic lattice before the runs are terminated. Our theory involves simple matrix algebra, which can be straightforwardly implemented numerically. The key assumption is that the track is periodic, that is the set of states available to the system and the rates that connect them do not depend on the motor location. This is physically a reasonable assumption for processive motors of the kinesin, dynein, or myosin families, which step along cytoplasmic tracks. We have also assumed that the track is infinitely long, which allows us to avoid introducing boundary effects. As an illustration, we applied the theory to study myosin VI processivity, and we fit the parameters to recover the experimental data (27, 54). We started from the experimental evidence on myosin VI motility provided by Yanagida and coworkers, and constructed a kinetic model in which both hand-over-hand and inchworm-like steps are considered. Our model shows that ADP-release gating and mechanical gating account for the average run length, velocity, and probability of forward stepping. The importance of ADP gating was highlighted by Elting et al. (27) and by Dunn et al. (49) on the basis of kinetic models that did not include backward stepping. Our model also reproduces the finding that backward steps are shorter than forward steps (52, 60), and agrees with a recent analysis suggesting that backward steps are mainly spontaneous (54). In addition, we make the important prediction that at high ATP concentrations, stomps break the tight mechano-chemical coupling of myosin VI by wasting approximately 13% of the hydrolyzed ATP molecules.

The theoretical framework propose could be improved in at least two aspects. First, he physical velocity is better defined as ⟨*L/τ* ⟩ instead of ⟨*L* ⟩*/* ⟨*τ* ⟩ We obtained the former with kinetic Monte Carlo (KMC) simulations (62), and we show that it differs from the latter (see Fig. S9d). Although both ⟨*L* ⟩ and ⟨*τ* ⟨ can be measured, thereby making ⟨ *L*⟩ */*⟨ *τ* ⟩ amenable to experimental testing, it would be informative to obtain *L/τ* for each stepping trajectory of the motor in order to measure the velocity distribution. It was predicted earlier with a simpler model that *P* (*L/τ*) is not Gaussian (38), and that aspect also holds for our model (see Fig. S5d and Fig. S6d). Second, in our model we overestimate the number of states. For instance, when one head detaches from the D or C state, the resulting 1HB states are identical to each other. However, in order to keep track of the stepping mechanism we “split” these states into two, one coming from the D state, another from the C state. This means that, as an example, a forward inchworm-like step may connect a 1HB C state with a 2HB D state, but the reverse transition is not possible because the 1HB state accessed from the 2HB D state would be listed as D, not C, although they are the same *physical* state. It would be instructive to develop the theoretical framework further in order to produce a solution to this subtle issue.

Finally, we mention that it is possible to obtain the results here using an alternative approach. Following (28, 63, 64), in the matrix 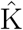 of Eq. 11 we may replace all the transitions towards the absorbing state with transitions back to the states that populate the initial probability distribution. This procedure replenishes the system with all of the trajectories absorbed in the detached state by restarting them with the appropriate probability distributions. The trajectories queued serially generate a stationary probability distribution 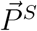 of populating the *N* states available to the system. From these stationary probabilities it is possible to obtain the fluxes for forward and backward stepping, and the flux to detachment, from which the average run length can be calculated (28). We verified numerically that the averages obtained with our approach and with this steady-state method are in excellent agreement.

## Methods

### Implementation of the model

The model for myosin VI was implemented in a Jupyter notebook (65), version 5.4.0. Functions from *NumPy* (66) were used to perform the matrix operations. The kinetic Monte Carlo code was also implemented in the same notebook, using the rejection-free scheme (62). Fitting was performed using the function *curve fit* of *SciPy* (67), and plots were made using *matplotlib* (68).

### Parameter optimization

We digitized the data points (ATP concentrations, values of the observables and error bars) used for learning the parameters of the model using *WebPlotDigitizer* (61) from Fig. S6c-d of Elting *et al.* (27) *and* Fig. 3B of Ikezaki *et al.* (54). Let [ATP]_*i*_ be the digitized ATP concentrations, and let *y*_exp_([ATP]_*i*_) be the digitized value of observable *y* at ATP concentration [ATP]_*i*_, with error given by *σ*_*y*,exp_([ATP]_*i*_); *y*_the_([ATP]_*i*_) is the value of *y* obtained using our model. The residual error is,

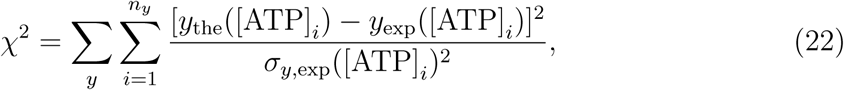

where the sum over *y* refers to different observables, and *n*_*y*_ is the number of ATP concentrations considered for *y*. The fit is performed by finding the set of parameters that minimize *χ*^2^ – we used the function *minimize* from *SciPy* (67). The parameters were all initialized at 1, and during the optimization, they were allowed to vary between *10*^*-*2^ and 10^2^.

## Supporting information

Details of the theory and simulations

## Acknowledgments

This work was supported by the founding from the National Science Foundation (CHE 19-00093), the Welch Foundation through the Collie-Welch chair (F-0019), the Center for Engineering MechanoBiology (CEMB), an NSF Science and Technology Center, under grant agreement CMMI: 15-48571, the National Institute of Health (NIH-5T32AR053461-to MC, and NIH grant R35GM118139 to YEG.

