## Supplementary material for "Processivity and Velocity for Motors Stepping on Periodic Tracks": Details of the theory and simulations

June 26, 2019

| | |
|---|---|
| $\vec{P}(x)$ | Probability of being in the $N$ accessible states after $x$ transitions have occurred |
| $\vec{P}(t)$ | Probability of being in the $N$ accessible states at time $t$ |
| $\hat{\mathrm{S}}$ | Changes in the chemical state at a fixed site |
| $\hat{\mathrm{F}}$ | Forward steps |
| $\hat{\mathrm{B}}$ | Backward steps |
| $\hat{\mathrm{M}}$ | Sum of all transitions excluding detachment (e.g. $\hat{\mathrm{M}} = \hat{\mathrm{S}} + \hat{\mathrm{F}} + \hat{\mathrm{B}}$) |
| $\hat{\mathrm{Q}}$ | Any set of transitions associated with the occurrence of an observable event |
| $\hat{\mathrm{K}}$ | Rate matrix |
| $\hat{\mathrm{S}}_{\mathrm{D}}$ | Changes in chemical state at a fixed site for conformations with distal heads (type $D$) |
| $\hat{\mathrm{S}}_{\mathrm{C}}$ | Changes in chemical state at a fixed site for conformations with close heads (type $C$) |
| $\hat{\mathrm{F}}_{\mathrm{HoH}}$ | Forward hand-over-hand step |
| $\hat{\mathrm{B}}_{\mathrm{HoH}}$ | Backward hand-over-hand step |
| $\hat{\mathrm{F}}_{\mathrm{Iw}}$ | Forward inchworm-like step |
| $\hat{\mathrm{B}}_{\mathrm{Iw}}$ | Backward inchworm-like step |
| $\hat{\mathrm{F}}_{\mathrm{IwD}}$ | Forward inchworm-like step from conformations $D$ to $C$ |
| $\hat{\mathrm{F}}_{\mathrm{IwC}}$ | Forward inchworm-like step from conformations $C$ to $D$ |
| $\hat{\mathrm{B}}_{\mathrm{IwD}}$ | Backward inchworm-like step from conformations $D$ to $C$ |
| $\hat{\mathrm{B}}_{\mathrm{IwC}}$ | Backward inchworm-like step from conformations $C$ to $D$ |
| $P(x)$ | Probability of being bound after $x$ transitions ($= \vec{1}^{\intercal} \cdot \vec{P}(x)$) |
| $\mathbb{P}(n)$ | Probability distribution that detachment occurs after $n$ steps |

Table S1: List of the symbols. The non-zero elements of each matrix or vector correspond to the transitions described in the second column.

| | |
|---|---|
| ADP release rate | |
| $k_{D,off}$ | actin-bound head without inter-head tension |
| $k'_{D,off}$ | head detached from actin |
| $k^{TH}_{D,off}$ | TH |
| $k^{LH}_{D,off}$ | LH |
| ADP binding rate | |
| $k_{D,on}$ | actin-bound head without inter-head tension |
| $k'_{D,on}$ | head detached from actin |
| $k^{TH}_{D,on}$ | TH |
| $k^{LH}_{D,on}$ | LH |
| ADP dissociation constant | |
| $K_D$ | actin-bound head without inter-head tension ($= k_{D,off}/k_{D,on}$) |
| $K'_D$ | head detached from actin ($= k'_{D,off}/k'_{D,on}$) |
| $K^{TH}_D$ | TH ($= k^{TH}_{D,off}/k^{TH}_{D,on}$) |
| $K^{LH}_D$ | LH ($= k^{LH}_{D,off}/k^{LH}_{D,on}$) |
| Phosphate release rate | |
| $k'_{P_i,off}$ | head detached from actin |
| ATP binding rate | |
| $k_{T,on}$ | actin-bound head without inter-head tension |
| $k'_{T,on}$ | head detached from actin |
| $k^{TH}_{T,on}$ | TH |
| $k^{LH}_{T,on}$ | LH |
| Spontaneous detachment | |
| $k_{M,off}$ | rate for spontaneous detachment from actin in the apo state in the absence of inter-head tension |
| $k_{MD,off}$ | rate for spontaneous detachment from actin in the ADP-bound state in the absence of inter-head tension |
| $s$ | enhancement of the rate of spontaneous detachment for myosin under inter-head tension (the rates become $sk_{M,off}$ and $sk_{MD,off}$) |
| Actin binding rate | |
| $l_X$ | rate for binding actin as a leading head in state D (X = M·T, M, or M·D) |
| $c_X$ | rate for binding actin at a site adjacent to the bound head (state C) (X = M·T, M, or M·D) |
| $t_X$ | rate for binding actin as a trailing head in state D (X = M·T, M, or M·D) |
| $k_{reb}$ | rate for actin binding for a detached head in the M·T state ($l_{MT} + c_{MT} + t_{MT} = k_{reb}$) |
| $r$ | enhancement factor when the detached head is in apo or ADP-bound state ($l_{MD} + c_{MD} + t_{MD} = rk_{reb}$) |
| $\theta$ | thermodynamic constraint relating the rate of actin binding in the apo and in the ADP-bound state ($l_M + c_M + t_M = r\theta k_{reb}, \theta = l_M/l_{MD} = c_M/c_{MD} = t_M/t_{MD}$) |
| Gating parameters | |
| $g_{ADP}$ | ADP-release gating ($g_{ADP} = k^{TH}_{D,off}/k^{LH}_{D,off}$) |
| $g_{ATP}$ | ATP-binding gating ($g_{ATP} = k^{TH}_{T,on}/k^{LH}_{T,on}$) |
| $g_{MEC}$ | mechanical gating ($g_{MEC} = l_{MT}/c_{MT} = c_{MT}/t_{MT} = c_{MD}/l_{MD} = c_{MD}/t_{MD} = c_M/l_M = c_M/t_M$) |

Table S2: List of the rates and related symbols.

### Derivation of the Main Results

In this first section we derive the key results of the paper. We use the following two identities (S1). First, if the moduli of all of the eigenvalues of a square matrix $\hat{\mathrm{X}}$ are $< 1$, the geometric series of $\hat{\mathrm{X}}$ is,

$$\sum_{n=0}^{\infty} \hat{\mathrm{X}}^n = (\hat{\mathrm{I}} - \hat{\mathrm{X}})^{-1} \tag{S1}$$

Second, if $\alpha$ is a scalar and $(\hat{\mathrm{I}} - \alpha\hat{\mathrm{X}})$ is invertible,

$$\frac{d}{d\alpha}(\hat{\mathrm{I}} - \alpha\hat{\mathrm{X}})^{-1}|_{\alpha=1} = (\hat{\mathrm{I}} - \hat{\mathrm{X}})^{-1}\hat{\mathrm{X}}(\hat{\mathrm{I}} - \hat{\mathrm{X}})^{-1}. \tag{S2}$$

**Derivation of Eq. 1.** Consider the periodic network in Fig. 1. Let $\vec{p}(l, x)$ be an $N$-dimensional vector describing the probability that after $x$ transitions the motor is in any of the $N$ accessible states at location $l$ along the track. Following the definitions of $\hat{\mathrm{S}}$, $\hat{\mathrm{F}}$, and $\hat{\mathrm{B}}$ provided in the main text, the probability $\vec{p}(l, x + 1)$ is,

$$\vec{p}(l, x + 1) = \hat{\mathrm{S}}\vec{p}(l, x) + \hat{\mathrm{F}}\vec{p}(l - 1, x) + \hat{\mathrm{B}}\vec{p}(l + 1, x), \tag{S3}$$

For a periodic track the matrices $\hat{\mathrm{F}}$, $\hat{\mathrm{B}}$, and $\hat{\mathrm{S}}$ are the same at all the sites along the filament. Therefore, we construct a vector $\vec{P}(x) = \sum_l \vec{p}(l, x)$, which contains the probability of being in any of the $N$ available states anywhere along the track after $x$ transitions. If the lattice is infinitely long, we obtain Eq. 1 by summing both the r.h.s and the l.h.s of Eq. S3 over $l$.

**The matrix $\hat{\mathrm{I}} - \hat{\mathrm{M}}$.** From Eq. 1, the probability of being bound to the track after $x$ transitions is $P(x) = \vec{1}^{\intercal}\hat{\mathrm{M}}^x\vec{P}(0)$, where $P(x)$ is a decreasing function of $x$ $[P(x + 1) \leq P(x)]$. The probability of detaching after exactly $x$ transitions is given by $P(x) - P(x + 1) \geq 0$, and is given by,

$$\begin{aligned} P(x) - P(x + 1) &= \vec{1}^{\intercal}\hat{\mathrm{M}}^x\vec{P}(0) - \vec{1}^{\intercal}\hat{\mathrm{M}}^{x+1}\vec{P}(0) = \\ &= \vec{1}^{\intercal}(\hat{\mathrm{I}} - \hat{\mathrm{M}})\vec{P}(x). \end{aligned} \tag{S4}$$

Eq. S4 indicates that multiplying by $\hat{\mathrm{I}} - \hat{\mathrm{M}}$ enables determination of the probability that the $(x + 1)$ transition represents dissociation from the track.

**Derivation of Eq. 2 and Eq. 4.** The probability, $\mathbb{P}(0)$, of detaching before the motor completes any forward or backward step is given by,

$$\begin{aligned} \mathbb{P}(0) &= \vec{1}^{\intercal}(\hat{\mathrm{I}} - \hat{\mathrm{M}})(\hat{\mathrm{I}} + \hat{\mathrm{S}} + \hat{\mathrm{S}}^2 + ...)\vec{P}(0) \\ &= \vec{1}^{\intercal}(\hat{\mathrm{I}} - \hat{\mathrm{M}})(\hat{\mathrm{I}} - \hat{\mathrm{S}})^{-1}\vec{P}(0). \end{aligned} \tag{S5}$$

We justify this expression with the following arguments. By multiplying $\vec{P}(0)$ by $\hat{\mathrm{S}}^x$ we populate the states visited by the system after $x$ transitions, none of which constitutes a step. Of course, the motor could detach, therefore $\vec{1}^\intercal \hat{\mathrm{S}}^x \vec{P}(0) \leq 1$. Summing $\hat{\mathrm{S}}^x$ over $x$, and using Eq. S1 we obtain, $\sum_{x=0}^{\infty} \hat{\mathrm{S}}^x = (\hat{\mathrm{I}} - \hat{\mathrm{S}})^{-1}$. Therefore, the term $(\hat{\mathrm{I}} - \hat{\mathrm{S}})^{-1}$ includes all of the pathways made of any number of transitions occurring at a fixed site along the track. Next, we need to ensure that the motor detaches before any step (or any further transition) takes place. This is achieved by multiplying by $\hat{\mathrm{I}} - \hat{\mathrm{M}}$, as discussed in the previous section of the SI. In summary, the matrix $(\hat{\mathrm{I}} - \hat{\mathrm{M}})(\hat{\mathrm{I}} - \hat{\mathrm{S}})^{-1}$ accounts for all the pathways in which the motor changes its state at a fixed location along the filament $[(\hat{\mathrm{I}} - \hat{\mathrm{S}})^{-1}]$ and then dissociates $[(\hat{\mathrm{I}} - \hat{\mathrm{M}})]$.

Analogously, we can write $\mathbb{P}(1)$, the probability of detaching after one step,

$$\mathbb{P}(1) = \vec{1}^\intercal (\hat{\mathrm{I}} - \hat{\mathrm{M}})(\hat{\mathrm{I}} - \hat{\mathrm{S}})^{-1}(\hat{\mathrm{F}} + \hat{\mathrm{B}})(\hat{\mathrm{I}} - \hat{\mathrm{S}})^{-1}\vec{P}(0). \tag{S6}$$

In this case we account for all possible trajectories in which any number of transitions are made before a step $[(\hat{\mathrm{I}} - \hat{\mathrm{S}})^{-1}]$, then the motor moves either forward or backward $(\hat{\mathrm{F}} + \hat{\mathrm{B}})$. At the new filament site, $l \pm 1$, the motor changes its state without moving along the track $[(\hat{\mathrm{I}} - \hat{\mathrm{S}})^{-1}]$, and then dissociates $[(\hat{\mathrm{I}} - \hat{\mathrm{M}})]$.

It follows from Eq. (S6) that the probability of detaching after exactly $n$ steps is,

$$\mathbb{P}(n) = \vec{1}^\intercal \hat{\mathrm{P}}(n)\vec{P}(0) = \vec{1}^\intercal \hat{\mathrm{P}}_{\mathrm{out}}(\hat{\mathrm{P}}_{\mathrm{step}})^n \vec{P}(0), \tag{S7}$$

where $\hat{\mathrm{P}}_{\mathrm{out}} = (\hat{\mathrm{I}} - \hat{\mathrm{M}})(\hat{\mathrm{I}} - \hat{\mathrm{S}})^{-1}$ and $\hat{\mathrm{P}}_{\mathrm{step}} = (\hat{\mathrm{F}} + \hat{\mathrm{B}})(\hat{\mathrm{I}} - \hat{\mathrm{S}})^{-1}$. We note that $\hat{\mathrm{I}} - \hat{\mathrm{P}}_{\mathrm{step}} = \hat{\mathrm{P}}_{\mathrm{out}}$, which can be straightforwardly verified by replacing $\hat{\mathrm{I}} = (\hat{\mathrm{I}} - \hat{\mathrm{S}})(\hat{\mathrm{I}} - \hat{\mathrm{S}})^{-1}$ and carrying out the algebra. The identity $\hat{\mathrm{I}} - \hat{\mathrm{P}}_{\mathrm{step}} = \hat{\mathrm{P}}_{\mathrm{out}}$ also ensures that $\sum_n \mathbb{P}(n) = \vec{1}^\intercal \vec{P}(0) = 1$, because,

$$\sum_n \hat{\mathrm{P}}_{\mathrm{out}}(\hat{\mathrm{P}}_{\mathrm{step}})^n = \hat{\mathrm{P}}_{\mathrm{out}}(\hat{\mathrm{I}} - \hat{\mathrm{P}}_{\mathrm{step}})^{-1} = \hat{\mathrm{I}}. \tag{S8}$$

Equation 2 is derived by substituting $(\hat{\mathrm{I}} - \hat{\mathrm{P}}_{\mathrm{step}})$ for $\hat{\mathrm{P}}_{\mathrm{out}}$ in Eq. S7.

The average number of steps completed before detachment is given by,

$$\langle n \rangle = \sum_{n=0}^{\infty} n\mathbb{P}(n) = \vec{1}^\intercal \hat{\mathrm{P}}_{\mathrm{out}} \sum_{n=0}^{\infty} n(\hat{\mathrm{P}}_{\mathrm{step}})^n \vec{P}(0), \tag{S9}$$

The right hand side of Eq. (S9) may be written as,

$$\langle n \rangle = \vec{1}^\intercal \hat{\mathrm{P}}_{\mathrm{out}} \lim_{\alpha \to 1} \alpha \frac{d}{d\alpha} \sum_{n=0}^{\infty} \alpha^n (\hat{\mathrm{P}}_{\mathrm{step}})^n \vec{P}(0), \tag{S10}$$

where the limit is taken from below, so that the moduli of all eigenvalues of the matrix $\alpha\hat{\mathrm{P}}_{\mathrm{step}}$ are $< 1$. Using the two identities in Eq. S1 and Eq. S2, we obtain Eq. 4.

**Derivation of Eq. 5 and Eq. 7.** Following the same reasoning, the probability of detaching without taking any forward step is,

$$\begin{aligned}\mathbb{P}_F(0) &= \vec{1}^\intercal(\hat{\mathrm{I}} - \hat{\mathrm{M}})(\hat{\mathrm{I}} + \hat{\mathrm{S}} + \hat{\mathrm{B}} + \hat{\mathrm{S}}\hat{\mathrm{B}} + \hat{\mathrm{B}}\hat{\mathrm{S}} + ...)\vec{P}(0) \\ &= \vec{1}^\intercal(\hat{\mathrm{I}} - \hat{\mathrm{M}})[\hat{\mathrm{I}} - (\hat{\mathrm{S}} + \hat{\mathrm{B}})]^{-1}\vec{P}(0),\end{aligned} \tag{S11}$$

Here, the matrix $[\hat{\mathrm{I}} - (\hat{\mathrm{S}} + \hat{\mathrm{B}})]^{-1}$ accounts for all the pathways that include both backward steps and transitions at a fix location along the filament, and $(\hat{\mathrm{I}} - \hat{\mathrm{M}})$ allows us to extract the probability of dissociating from the track. Similarly, the probability of detaching after one forward step is,

$$\mathbb{P}_F(1) = \vec{1}^\intercal(\hat{\mathrm{I}} - \hat{\mathrm{M}})[\hat{\mathrm{I}} - (\hat{\mathrm{S}} + \hat{\mathrm{B}})]^{-1}\hat{\mathrm{F}}[\hat{\mathrm{I}} - (\hat{\mathrm{S}} + \hat{\mathrm{B}})]^{-1}\vec{P}(0). \tag{S12}$$

Generalizing as we did before, if we define $\hat{\mathrm{P}}_{\mathrm{F,out}} = (\hat{\mathrm{I}} - \hat{\mathrm{M}})[\hat{\mathrm{I}} - (\hat{\mathrm{S}} + \hat{\mathrm{B}})]^{-1}$ , and $\hat{\mathrm{P}}_{\mathrm{F}} = \hat{\mathrm{F}}[\hat{\mathrm{I}} - (\hat{\mathrm{S}} + \hat{\mathrm{B}})]^{-1}$, we get that the probability of taking $f$ forward steps is,

$$\mathbb{P}_F(f) = \vec{1}^\intercal(\hat{\mathrm{I}} - \hat{\mathrm{P}}_{\mathrm{F}})(\hat{\mathrm{P}}_{\mathrm{F}})^f\vec{P}(0), \tag{S13}$$

where we used the identity $\hat{\mathrm{P}}_{\mathrm{F,out}} = \hat{\mathrm{I}} - \hat{\mathrm{P}}_{\mathrm{F}}$, which can be proven by direct substitution, and ensures that $\sum_{f=0}^{\infty}\mathbb{P}_F(f) = 1$. The average number of forward steps is given by,

$$\langle f \rangle = \sum_{f=0}^{\infty} f\mathbb{P}_F(f), \tag{S14}$$

and one gets to Eq. 5 by following the same steps that lead to $\langle n \rangle$.

Analogously, the probability of dissociating from the track after $b$ backward steps have occurred is,

$$\mathbb{P}_B(b) = \vec{1}^\intercal(\hat{\mathrm{I}} - \hat{\mathrm{P}}_{\mathrm{B}})(\hat{\mathrm{P}}_{\mathrm{B}})^b\vec{P}(0), \tag{S15}$$

where $\hat{\mathrm{P}}_{\mathrm{B}} = \hat{\mathrm{B}}[\hat{\mathrm{I}} - (\hat{\mathrm{S}} + \hat{\mathrm{F}})]^{-1}$. In order to derive Eq. 7, we compute,

$$\langle b \rangle = \sum_{b=0}^{\infty} b\mathbb{P}_B(b). \tag{S16}$$

**Derivation of Eq. 9 and Eq. 10.** Equations 9 and 10 may be obtained by recognizing that $\langle n \rangle = \langle f \rangle + \langle b \rangle$. This identity must hold regardless of the initial probability distribution $\vec{P}(0)$. Therefore, we must show that,

$$\begin{aligned}&\hat{\mathrm{P}}_{\mathrm{step}}(\hat{\mathrm{I}} - \hat{\mathrm{P}}_{\mathrm{step}})^{-1} = \\ &\qquad \hat{\mathrm{P}}_{\mathrm{F}}(\hat{\mathrm{I}} - \hat{\mathrm{P}}_{\mathrm{F}})^{-1} + \hat{\mathrm{P}}_{\mathrm{B}}(\hat{\mathrm{I}} - \hat{\mathrm{P}}_{\mathrm{B}})^{-1}.\end{aligned} \tag{S17}$$

On the l.h.s. of Eq. S17, we note that $(\hat{\rm I}-\hat{\rm P}_{\rm step})^{-1}=(\hat{\rm I}-\hat{\rm S})(\hat{\rm I}-\hat{\rm M})^{-1}$, and therefore,

$$\hat{\rm P}_{\rm step}(\hat{\rm I}-\hat{\rm P}_{\rm step})=(\hat{\rm F}+\hat{\rm B})(\hat{\rm I}-\hat{\rm M})^{-1}.$$

Equation S17 is verified by applying the same transformation to the r.h.s.

#### Variable Step Size

**Derivation of Eq. 14.** To account for both the inchworm and hand-over-hand stepping we consider a periodic lattice with two sets of intertwined sites along the filament.The D states are labeled by integers $..., l-1, l, l+1, ...$ and the sites identified by half-integers $..., l-1/2, l+1/2, ...$ correspond to C states (see Fig. 3). The $N$-dimensional vectors $\vec{p}_{\rm D}(x,l)$ and $\vec{p}_{\rm C}(x,l+1/2)$ define the probabilities of being in any of the $N_{\rm D}$ and $N_{\rm C}$ conformations of type D and C, respectively. Note that $\hat{\rm S}_{\rm D}$, $\hat{\rm F}_{\rm IwD}$, and $\hat{\rm B}_{\rm IwD}$ operate only on the D states. Hand-over-hand steps cannot occur when the heads are close (C states), and $\hat{\rm S}_{\rm C}$, $\hat{\rm F}_{\rm IwC}$, and $\hat{\rm B}_{\rm IwC}$ affect only on the C states. Therefore,

$$\begin{array}{ll} \hat{\rm S}_{\rm C}\vec{p}_{\rm D}(x,l)=0 & \hat{\rm S}_{\rm D}\vec{p}_{\rm C}(x,l+\frac{1}{2})=0 \\ \hat{\rm F}_{\rm HoH}\vec{p}_{\rm C}(x,l+\frac{1}{2})=0 & \hat{\rm B}_{\rm HoH}\vec{p}_{\rm C}(x,l+\frac{1}{2})=0 \\ \hat{\rm F}_{\rm IwC}\vec{p}_{\rm D}(x,l)=0 & \hat{\rm B}_{\rm IwC}\vec{p}_{\rm D}(x,l)=0 \\ \hat{\rm F}_{\rm IwD}\vec{p}_{\rm C}(x,l+\frac{1}{2})=0 & \hat{\rm B}_{\rm IwD}\vec{p}_{\rm C}(x,l+\frac{1}{2})=0. \end{array} \tag{S18}$$

The Markov chains that governs the evolution of the D and C states are,

$$\begin{aligned} \vec{p}_{\rm D}(x+1,l) &= \hat{\rm S}_{\rm D}\vec{p}_{\rm D}(x,l)+ \\ &+\hat{\rm F}_{\rm HoH}\vec{p}_{\rm D}(x,l-1)+\hat{\rm B}_{\rm HoH}\vec{p}_{\rm D}(x,l+1)+ \\ &+\hat{\rm F}_{\rm IwC}\vec{p}_{\rm C}(x,l-\frac{1}{2})+\hat{\rm B}_{\rm IwC}\vec{p}_{\rm C}(x,l+\frac{1}{2}) \\ \vec{p}_{\rm C}(x+1,l+\frac{1}{2}) &= \hat{\rm S}_{\rm C}\vec{p}_{\rm C}(x,l+\frac{1}{2})+ \\ &+\hat{\rm F}_{\rm IwD}\vec{p}_{\rm D}(x,l)+\hat{\rm B}_{\rm IwD}\vec{p}_{\rm D}(x,l+1) \end{aligned} \tag{S19}$$

We construct a vector $\vec{P}(x)=\sum_l[\vec{p}_{\rm D}(x,l)+\vec{p}_{\rm C}(x,l+1/2)]$ by assuming that the filament is infinitely long. Because all the matrices are identical at each lattice site, we can sum over $l$ the r.h.s and the l.h.s of Eq. S19, and using Eq. S18 we obtain Eq. 14.

**Average Number of Steps.** The distributions and average number of steps can be obtained as shown before. If we define $\hat{\rm F}=\hat{\rm F}_{\rm HoH}+\hat{\rm F}_{\rm Iw}$ and $\hat{\rm B}=\hat{\rm B}_{\rm HoH}+\hat{\rm B}_{\rm Iw}$, $\langle n\rangle$ is the same as in Eq. 4, the average number of forward hand-over-hand steps is the same as Eq. 5, with $\hat{\rm P}_{\rm F}=\hat{\rm F}_{\rm HoH}[\hat{\rm I}-(\hat{\rm S}+\hat{\rm F}_{\rm Iw}+\hat{\rm B})]^{-1}$, whereas in order to obtain the average number of forward inchworm-like steps it is necessary to replace $\hat{\rm P}_{\rm F}=\hat{\rm F}_{\rm Iw}[\hat{\rm I}-(\hat{\rm S}+\hat{\rm F}_{\rm HoH}+\hat{\rm B})]^{-1}$ in Eq. 5. The average backward steps for the two stepping modes can be constructed accordingly.

**Probability, Average Run Length, and Velocity.** The probability of taking a forward/backward hand-over-hand/inchworm-like step is given by the ratio between the average number of times a specific type of step occurs and the total number of steps. For this to hold, $\langle n \rangle$ must be equal to the sum of the four stepping modes $\langle f_{\mathrm{HoH}} \rangle$, $\langle f_{\mathrm{Iw}} \rangle$, $\langle b_{\mathrm{HoH}} \rangle$, and $\langle b_{\mathrm{Iw}} \rangle$. This can be proven following the strategy used to prove Eq. S17.

For the average run length, we consider Eq. 16, in which we assumed that inchworm steps are half of the size of the hand-over-hand step. In general, one could distinguish two types of inchworm steps, those going from states D to C, and the transitions from C to D. Depending on the location of the probe on the motor the step-sizes might not be identical.

In order to obtain the average velocity when inchworm steps are included, one must redefine the matrix $\hat{\mathrm{K}}$ in Eq. 11 following the same strategy discussed for the Markov chain.

**Derivation of Eq. 17 and Eq. 18.** Let $\hat{\mathrm{Q}}$ be the matrix containing all the transitions at which a monitored event , such as ATP binding or backward stepping, occurs. We are interested in the probability distribution of $\mathbb{P}_{\mathrm{Q}}(q)$, where $q$ is the number of times the event takes place before detachment from the track. We follow the derivation of $\mathbb{P}(n)$ illustrated at the beginning of the Appendix. By defining $\hat{\mathrm{P}}_{\mathrm{Q}}$ as in Eq. 18 we obtain,

$$\mathbb{P}_{\mathrm{Q}}(q) = \vec{\mathrm{1}}^{\intercal}(\hat{\mathrm{I}} - \hat{\mathrm{P}}_{\mathrm{Q}})(\hat{\mathrm{P}}_{\mathrm{Q}})^{q}\vec{P}(0). \tag{S20}$$

Eq. 17 is the average $\langle q \rangle = \sum_q q\mathbb{P}_Q(q)$, and can be derived as done previously for the average number of steps.

#### Initial Condition

We assume that a myosin head detached from the filament exists in only three states: apo (M), ADP-bound (M·D), and in post-ATP-hydrolysis state (M·D·$\mathrm{P_i}$). Let the solution be such that there is no free phosphate in solution, and that ATP binding is irreversible. In this case, the transitions allowed are in Fig. S1. In a stationary state, the populations of M·D, M, and M·T are given by,

$$\begin{aligned}
\mathrm{P_{M\cdot D}} &= \frac{(\mathrm{k}'_{\mathrm{T,on}}[\mathrm{T}] + \mathrm{k}'_{\mathrm{D,on}}[\mathrm{D}])\mathrm{k}'_{\mathrm{P_i,off}}}{\Sigma} \\
\mathrm{P_{M}} &= \frac{\mathrm{k}'_{\mathrm{D,off}}\mathrm{k}'_{\mathrm{P_i,off}}}{\Sigma} \\
\mathrm{P_{M\cdot D\cdot P_i}} &= \frac{\mathrm{k}'_{\mathrm{T,on}}[\mathrm{T}]\mathrm{k}'_{\mathrm{D,off}}}{\Sigma}
\end{aligned} \tag{S21}$$

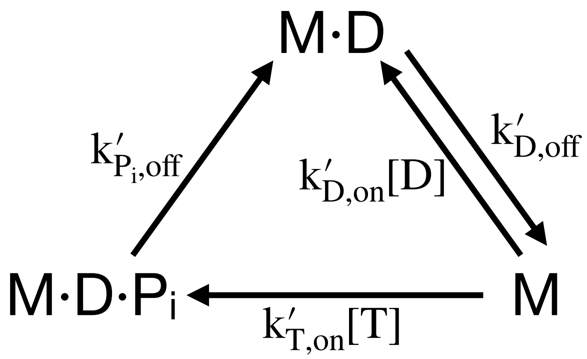


Figure S1: States available to the detached myosin head. It is assumed that the solution has no free $P_i$, and that ATP binding is irreversible. M·D is ADP-bound, M is apo, and M·D·$P_i$ is post-ATP-hydrolysis state, bound to ADP and $P_i$. The transition M·D → M is due to ADP release, ATP binding to M leads to M·D·$P_i$, and M·D·$P_i$ → M·D occurs upon release of phosphate. M·D·$P_i$ is a pre-power stroke state, M·D and M are post-power stroke states.

where $\Sigma = (k'_{T,on}[T] + k'_{D,on}[D])k'_{P_i,off} + k'_{D,off}k'_{P_i,off} + k'_{T,on}[T]k'_{D,off}$. In order to define the probability of binding to actin in a particular biochemical state with a given geometry we assume the following. (1) When the two heads are detached, they are independent from each other. (2) The lever arms of apo and ADP-bound myosin are at the post-stroke angle, in contrast to the orientation of the M·D·$P_i$ lever arm. If both of the bound heads are in their low-energy angle statess, binding is enhanced by a factor $g^2_{MEC}$. If only one head is in the low energy state the factor is $g_{MEC}$. It follows that the probability of binding to actin in conformation D is,

$$\begin{aligned}
\Sigma P_{AM\cdot D.AM\cdot D} &= {P_{M\cdot D}}^2 g_{MEC} + P_{M\cdot D}P_{M\cdot D\cdot P_i}g^2_{MEC} + \\
&+ P_{M\cdot D\cdot P_i}P_{M\cdot D} + P^2_{M\cdot D\cdot P_i}g_{MEC} \\
\Sigma P_{AM.AM\cdot D} &= P_M P_{M\cdot D} g_{MEC} + P_M P_{M\cdot D\cdot P_i} g^2_{MEC} \\
\Sigma P_{AM\cdot D.AM} &= P_{M\cdot D}P_M g_{MEC} + P_{M\cdot D\cdot P_i}P_M \\
\Sigma P_{AM.AM} &= {P_M}^2 g_{MEC},
\end{aligned} \tag{S22}$$

whereas for conformation C we obtain,

$$\begin{aligned}
\Sigma P_{AM\cdot D.AM\cdot D} &= {P_{M\cdot D}}^2 g^2_{MEC} + 2P_{M\cdot D}P_{M\cdot D\cdot P_i}g_{MEC} + \\
&+ P^2_{M\cdot D\cdot P_i}g^2_{MEC} \\
\Sigma P_{AM\cdot D.AM} &= 2P_{M\cdot D}P_M g^2_{MEC} + 2P_{M\cdot D\cdot P_i}P_M g_{MEC} \\
\Sigma P_{AM.AM} &= {P_M}^2 g^2_{MEC}.
\end{aligned} \tag{S23}$$

Here, $\Sigma$ is the sum of all the r.h.s. Note that in conformation C AM·D.AM is the same as AM.AM·D.

### Mechanical Gating

In Fig. S2 we illustrate graphically the effect of mechanical gating. As explained in the main text, the lever arm of an apo or ADP-bound myosin would preferentially orient forward (post-power-stroke conformation) in the absence of inter-molecular strain from its partner head. After ATP hydrolysis, the re-priming stroke orients the lever arm preferentially backward. The likelihood of binding to actin in a 2HB conformation is enhanced by a factor $g_{\mathrm{MEC}}$ for each lever arm in its favored conformation. A stepping head can bind actin in three locations (see Table S2), ahead of the bound motor (with rate $l_X$, where $X = \mathrm{M}$, M·T, or M·D, depending on the nucleotide bound to the stepping head), in trailing position (with rate $t_X$), or in the "center", that is adjacent to the bound motor (with rate $c_X$). The factor $r$ (see Table S2) is the enhancement in the rate of binding actin in the apo and ADP-bound states compared to the ATP-bound head. The factor $\theta$ comes from thermodynamic considerations, which we discuss in the next section.

### Thermodynamic Constraints

In Fig. S3 we show 4 cycles, which define multiple pathways between states. The free energy difference upon completing a cycle is equal to zero, which sets constraints between some of the rates. Working out the algebra, we find from Fig. S3 that,

$$
\begin{aligned}
\mathrm{K_D^{TH}} &= \mathrm{K_D^{LH}} = \mathrm{K_D} \\
\frac{t_{\mathrm{M}}}{t_{\mathrm{MD}}}\frac{\mathrm{k^{TH}_{MD,off}}}{\mathrm{k^{TH}_{M,off}}} &= \frac{\mathrm{K_D}}{\mathrm{K'_D}} \\
\frac{t_{\mathrm{M}}}{t_{\mathrm{MD}}}\frac{\mathrm{k^{TH}_{MD,off}}}{\mathrm{k^{TH}_{M,off}}} &= \frac{l_{\mathrm{M}}}{l_{\mathrm{MD}}}\frac{\mathrm{k^{LH}_{MD,off}}}{\mathrm{k^{LH}_{M,off}}},
\end{aligned}
\tag{S24}
$$

where $\mathrm{K_D^{TH}}$ ($\mathrm{K_D^{LH}}$) is the ADP dissociation constant for the TH (LH), and $\mathrm{K_D}$ is the ADP dissociation constant for the bound head in a 1HB state (see Table S2). The first equation states that there is only one ADP dissociation constant for actin-bound myosin. The second equation relates the ADP binding constant for an actin-bound ($\mathrm{K_D} = \mathrm{k_{D,off}}/\mathrm{k_{D,on}}$) and a dissociated head ($\mathrm{K'_D} = \mathrm{k'_{D,off}}/\mathrm{k'_{D,on}}$) to a function of the rates of detachment and rebinding in trailing position of the ADP-bound and apo myosin TH. The last identity relates the rates for spontaneous detachment in the LH and TH to the rates of binding actin in leading or trailing position.

The two pathways displayed in Fig. S4E-F connect the conformation AM·D.AM·D of the motor at two consecutive sites along the filament. Nevertheless, because no energy is expended in any of these transitions, the forward and backward pathways must occur with the same probability. In combination with Eq. S24, we obtain the following constraints on the rates,

$$\begin{aligned} \mathrm{k}^{\mathrm{TH}}_{\mathrm{MD,off}} l_{\mathrm{MD}} &= \mathrm{k}^{\mathrm{LH}}_{\mathrm{MD,off}} t_{\mathrm{MD}} \\ \mathrm{k}^{\mathrm{TH}}_{\mathrm{M,off}} l_{\mathrm{M}} &= \mathrm{k}^{\mathrm{LH}}_{\mathrm{M,off}} t_{\mathrm{M}}. \end{aligned} \tag{S25}$$

The interpretation of these constraints is clear: the probability of spontaneous forward and backward stepping must be the same. Note that the third identity in Eq. S24 follows from Eq. S25.

In conformation C, with the two myosin heads bound close to each other, the thermodynamic cycle in Fig. S4G sets the following relationship,

$$\frac{c_{\mathrm{M}}}{c_{\mathrm{MD}}} \frac{\mathrm{k}_{\mathrm{MD,off}}}{\mathrm{k}_{\mathrm{M,off}}} = \frac{\mathrm{K}_{\mathrm{D}}}{\mathrm{K}'_{\mathrm{D}}}. \tag{S26}$$

We implemented the thermodynamic constraints with the following considerations: we assume that the effect of inter-molecular force on the spontaneous detachment rate from the apo and ADP-bound state is the same, and we introduced a fitting parameter $s$ so that,

$$\begin{aligned} \mathrm{k}^{\mathrm{TH}}_{\mathrm{MD}} &= \mathrm{k}^{\mathrm{LH}}_{\mathrm{MD}} = s\mathrm{k}_{\mathrm{MD}} \\ \mathrm{k}^{\mathrm{TH}}_{\mathrm{M}} &= \mathrm{k}^{\mathrm{LH}}_{\mathrm{M}} = s\mathrm{k}_{\mathrm{D}}. \end{aligned} \tag{S27}$$

As a consequence, it follows from Eqs. S24-S26 that,

$$\begin{aligned} l_{\mathrm{MD}} &= t_{\mathrm{MD}} \\ l_{\mathrm{M}} &= t_{\mathrm{M}} \\ l_{\mathrm{M}} &= l_{\mathrm{MD}} \frac{\mathrm{k}_{\mathrm{M,off}}}{\mathrm{k}_{\mathrm{MD,off}}} \frac{\mathrm{K}_{\mathrm{D}}}{\mathrm{K}'_{\mathrm{D}}} = \theta l_{\mathrm{MD}} \\ c_{\mathrm{M}} &= c_{\mathrm{MD}} \frac{\mathrm{k}_{\mathrm{M,off}}}{\mathrm{k}_{\mathrm{MD,off}}} \frac{\mathrm{K}_{\mathrm{D}}}{\mathrm{K}'_{\mathrm{D}}} = \theta c_{\mathrm{MD}} \end{aligned} \tag{S28}$$

The factor $\theta$ is not a fitting parameter, and is obtained from bulk kinetic experiments (see Table 1).

### Distributions

So far, we considered only average properties. In this section we discuss the run-length, velocity, and dwell-time distributions for the model in Fig. 3 with the rates obtained after fitting our model to the data in (S2) and (S3) (see Table 1). Eq. 2 in the main text shows how to obtain the distribution of the number of steps taken by the motor before the end of the processive run. We can compare the distribution with the results of kinetic Monte Carlo (KMC) simulations. It is challenging to derive analytical expressions for other distributions, such as the run length, attachment time, velocity, and dwell time. Therefore we only show the results for these quantities from KMC simulations, which we fit to simple functions (see Fig. S5 for [ATP] = 1 mM and Fig. S6 for [ATP] = 100 $\mu$M).

The agreement between the analytical and numerical $\mathbb{P}(n)$ is excellent. Excluding the peaks at $L = 0$ and $t = 0$, the distributions for run length and run time are well described by a single exponentials. The velocity distribution is multimodal, as predicted for kinesin (S4): there is an extremely small population for $v < 0$, a peak at $v = 0$ referring to all the myosin molecules that detach without stepping, and a broader population for $v > 0$. The sub-population at positive velocities cannot be accurately described by a Gaussian distribution.

The distribution of dwell-times between two steps is frequently studied experimentally. We extract the dwell time distribution from the model at [ATP] = 1 mM and [ATP] = 100 $\mu$M from KMC simulations. We distinguish between the dwell time before a hand-over-hand forward or backward step, and an inchworm-like step in both directions (see Fig. S7 and S8). All of them are reasonably well fitted by a convolution of two exponentials.

### Alternate Definitions of Velocity

As explained in the main text, we adopted $\langle L\rangle/\langle\tau\rangle$ as a definition of velocity. This needs not to be the same as $\langle L/\tau\rangle$, which can be computed with KMC simulations. In Fig. S9 we show the comparison between the two. First, the number of steps, average run length, and average velocity ($\langle L\rangle/\langle\tau\rangle$) are the same in KMC and in the theory described in the paper (Figs. S9a-S9c). Fourth, we plot the relative error made using $\langle L\rangle/\langle\tau\rangle$ instead of $\langle L/\tau\rangle$ as a function of the average number of steps taken by the motor (Fig. S9d). The relative error between $\langle L/\tau\rangle$ and $\langle L\rangle/\langle\tau\rangle$ decreases as $N$ increases.

### Run Length in the Absence of ATP

In Fig. S10a we show that as [ATP] decreases, eventually the run length decreases as well – as expected, the value found at [ATP] = 0 is essentially zero ($|L| \approx 5 \times 10^{-11}$nm). As the concentration of ADP is varied (with [ATP] = 0), there is a weak dependence of the run length, which reaches a peak around $\approx 10 - 100\,\mu$M (Fig. S10b). The system is not in equilibrium because of the absorbing boundary condition, and the initial state might inject a small amount of energy which results in a non-zero (albeit small) average run length even in the absence of ATP. The calculations are performed using the theory described in the paper, and the equivalent steady-state method briefly mentioned in the Discussion in the main text.

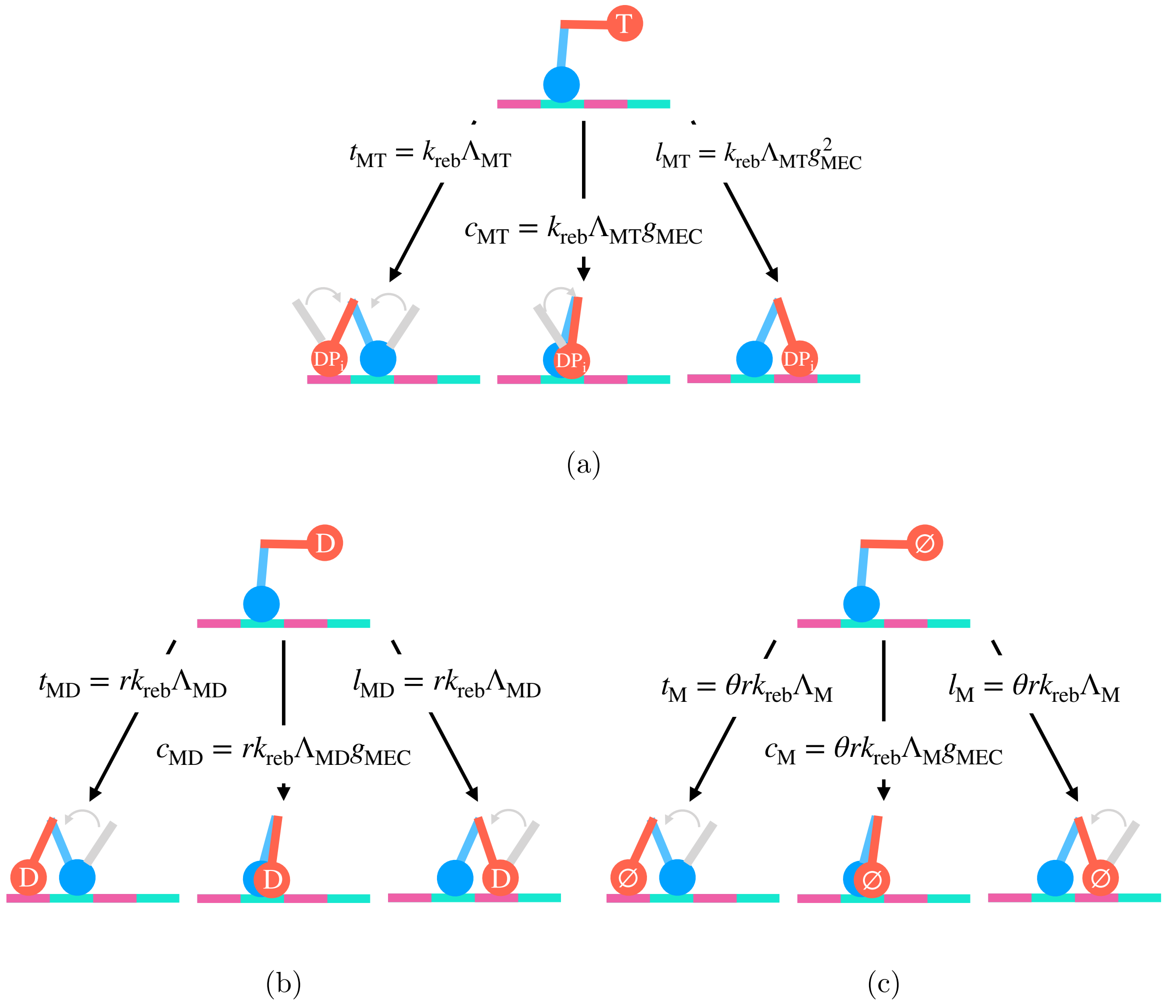


Figure S2: Implementation of the gating mechanism parameter. The stepping head is shown in red, the blue head is bound (whether it is ADP-bound or apo is irrelevant). The arrows show three binding options: trailing position, adjacent to the bound head, or distal to the bound head. The grey lever arms indicate the preferential orientation of a head if it were not under strain in a 2HB state. (a) Effect of mechanical gating on the completion of a ATP-initiated step that is initiated by ATP binding [$\Lambda_{\mathrm{MT}} = (g^2_{\mathrm{MEC}} + g_{\mathrm{MEC}} + 1)^{-1}$]. (b) Mechanical gating when ADP is bound to the stepping head (M·D state), with $\Lambda_{\mathrm{MD}} = (g_{\mathrm{MEC}} + 2)^{-1}$. (c) Effect of mechanical gating when the stepping motor is in the apo state [$\Lambda_{\mathrm{M}} = (g_{\mathrm{MEC}} + 2)^{-1}$]. The rates shown in the figures are defined in Eqs. 19-20 and in Table S2.

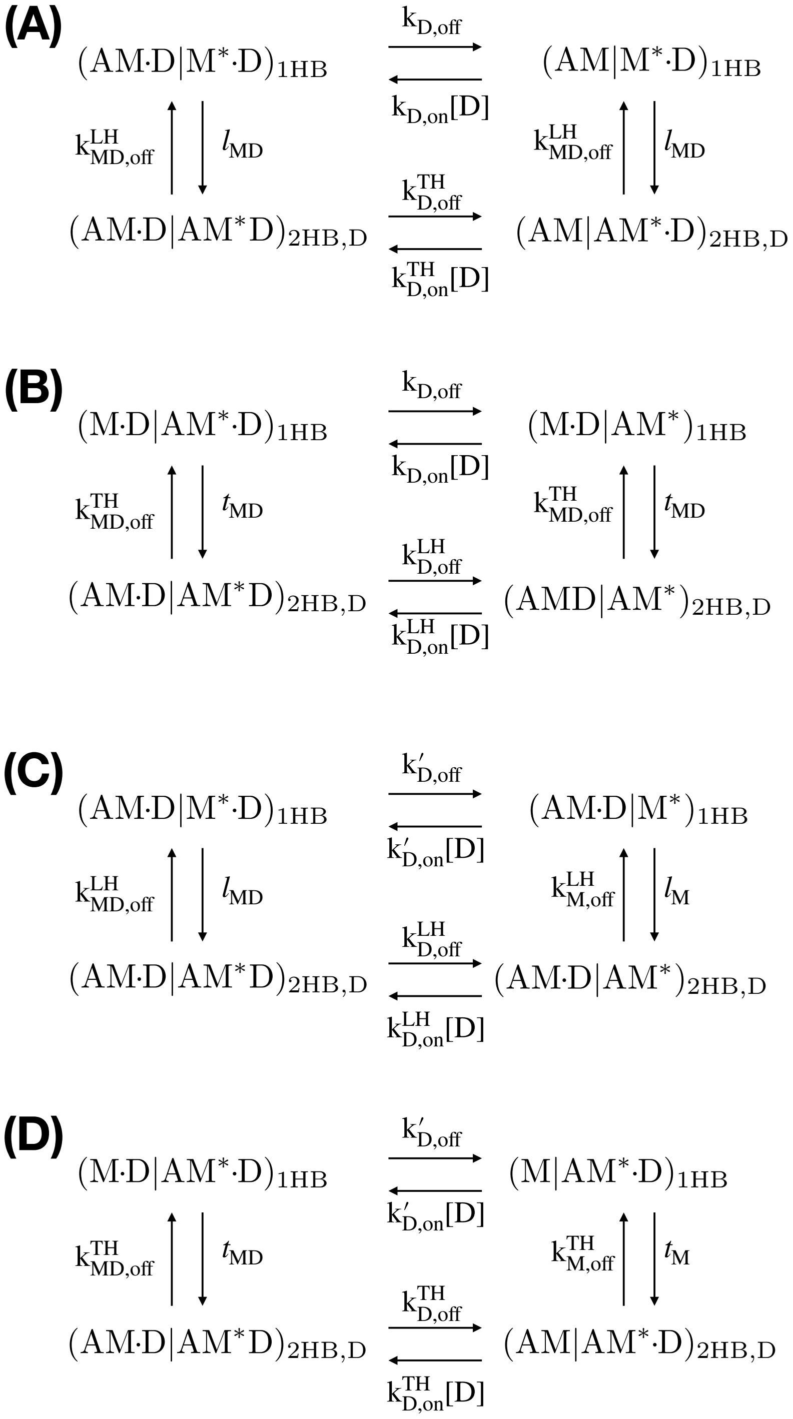


Figure S3: Thermodynamic constraints on the model. The kinetic schemes illustrate four cycles that sets constraints between the rates in the model. The symbol $(\mathrm{AM{\cdot}D|AM^*})_{\mathrm{2HB,D}}$ refers to a 2HB motor in conformation D. ADP is bound to the TH, the LH is in the apo state. One of the two heads is "labeled" with a star in order to distinguish between TH and LH..

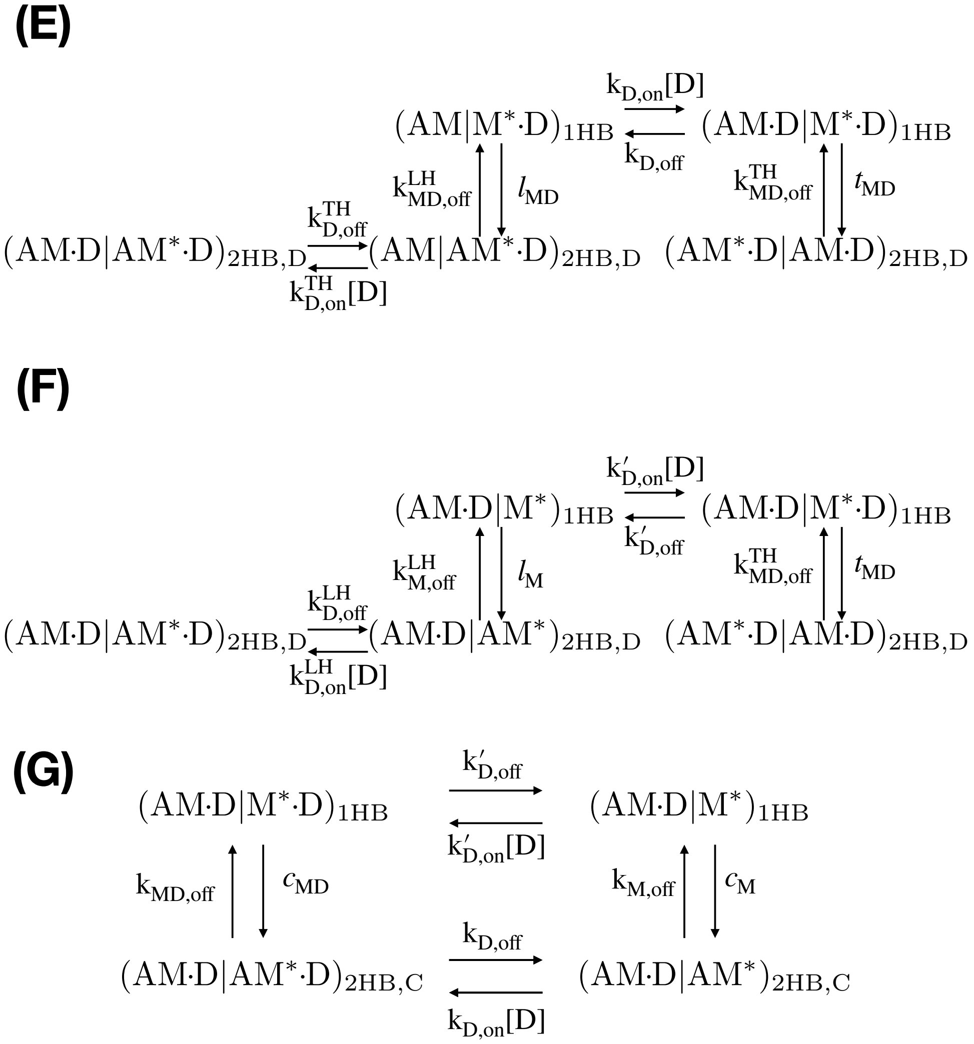


Figure S4: Thermodynamic constraints on the model. E and F show pathways for spontaneous forward and backward stepping in state D. G refers to myosin with the heads bound to adjacent actin sites (state C)

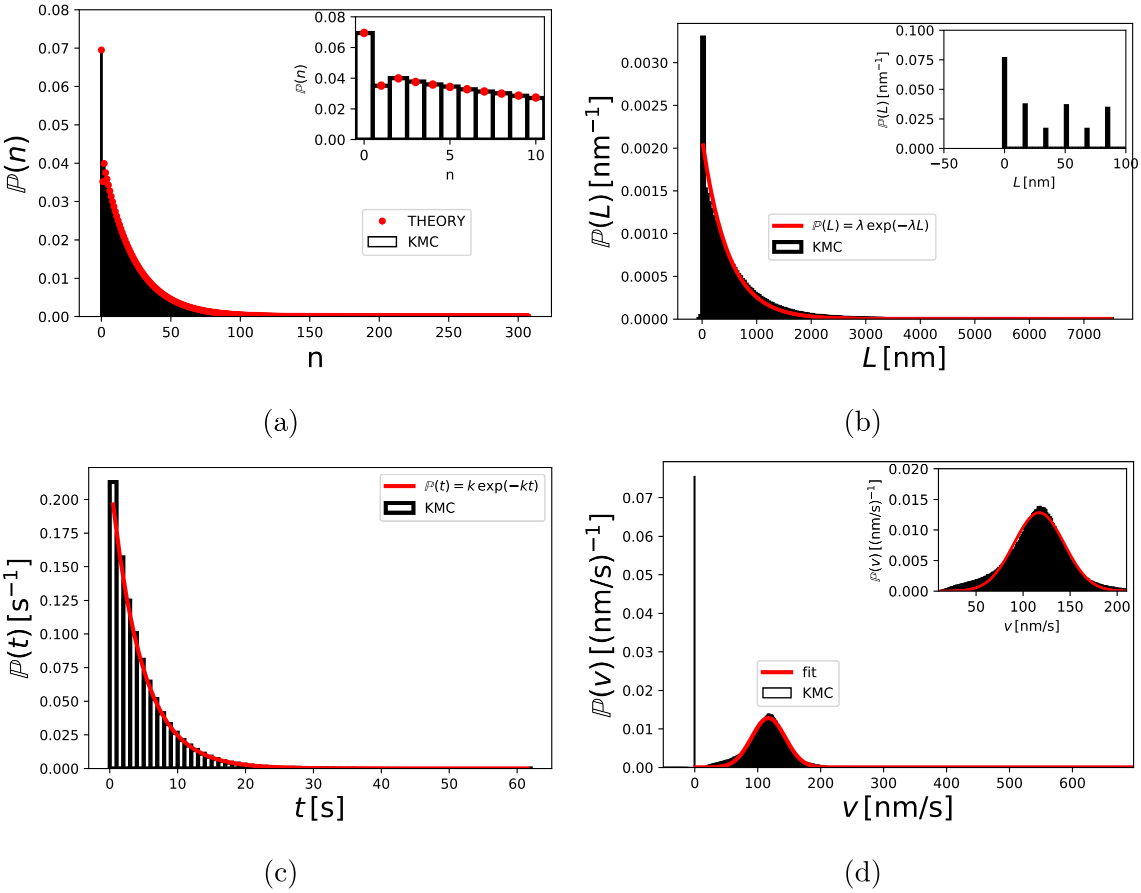


Figure S5: Distributions of various quantities at [ATP] = 1 mM. Histograms, in black, are from kinetic Monte Carlo (KMC) simulations. (a) Distribution of number of steps. The red dots are results calculated using Eq. 2. The inset highlights the initial region. (b) Run length distribution. The red line is from a fit to a single exponential ($\lambda e^{-\lambda L}$) excluding $L = 0$ [$\lambda = (2.11 \pm 0.05)10^{-3}\text{nm}^{-1}$]. The inset shows that $L$ takes on discrete values. (c) Run time distribution. The red line is a fit to a single exponential ($ke^{-kt}$) excluding $t = 0$ [$k = (0.2190 \pm 0.0007)\text{s}^{-1}$]. (d) Velocity distribution. The velocity is defined as $L/t$ for each trajectory. The red line is a fit of the positive velocities to a Gaussian distribution ($ae^{-(x-\mu)^2/(2\sigma^2)}$, with $a = (128.3 \pm 0.7)10^{-4}$, $\mu = 117.0 \pm 0.2$, $\sigma = 26.7 \pm 0.2$. All values are in nm/s). The inset highlights the peak of the velocity distribution.

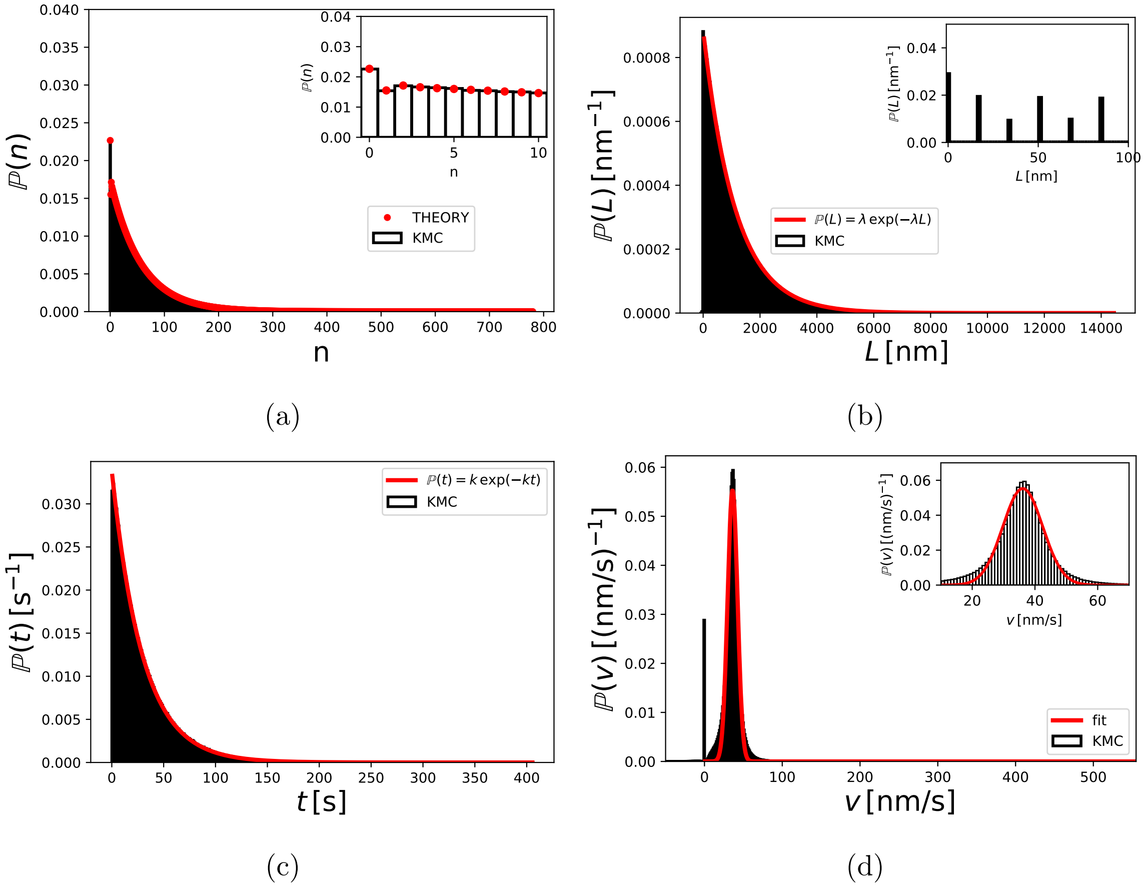


Figure S6: Distributions at [ATP] $= 100\,\mu$M. Same as the caption for Fig. S5. Fitted parameters: (b) run length, $\lambda = (8.859 \pm 0.009)10^{-4}\text{nm}^{-1}$, (c) run time, $k = (3.382 \pm 0.002)10^{-2}\text{s}^{-1}$, and (d) velocity, $a = (5.54 \pm 0.03)10^{-2}\text{nm/s}$, $\mu = (36.12 \pm 0.04)\text{nm/s}$, $\sigma = (6.36 \pm 0.05)\text{nm/s}$.

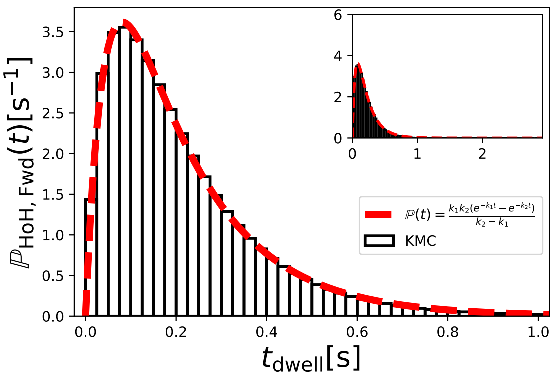


(a)

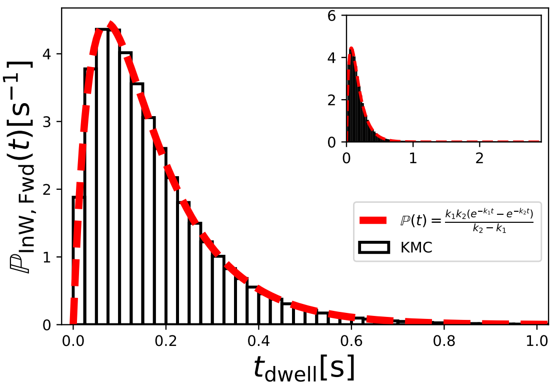


(b)

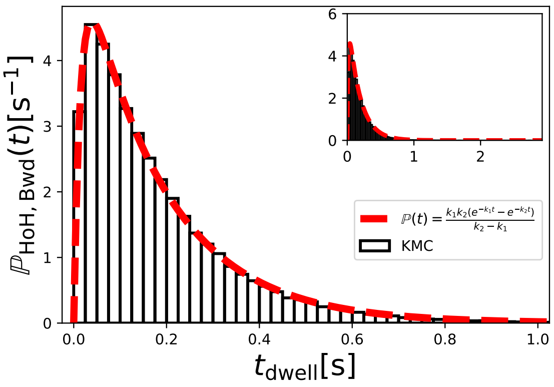


(c)

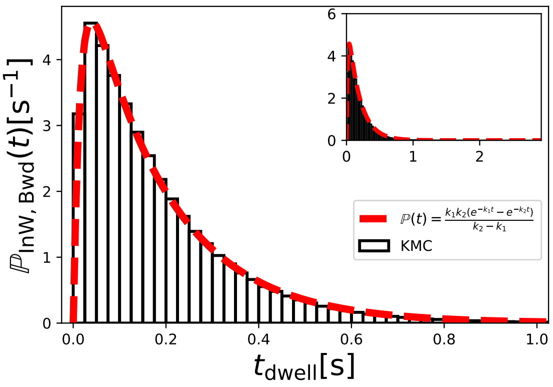


(d)

Figure S7: Dwell-time distributions at [ATP] = 1 mM. Histograms, in black, are from KMC simulations. Fits are in red. The insets show the entire distribution, the main figure focuses on the initial part. The fit is performed with a convolution between two exponentials $[k_1 k_2/(k_1 - k_2)(e^{-k_2 t} - e^{-k_1 t})]$. (a) Hand-over-hand forward steps $[k_1 = (22.7 \pm 0.2)\text{s}^{-1}$ and $k_2 = (5.78 \pm 0.02)\text{s}^{-1}]$. (b) Inchworm forward steps $[k_1 = (22.5 \pm 0.1)\text{s}^{-1}$ and $k_2 = (7.82 \pm 0.02)\text{s}^{-1}]$. (c) Hand-over-hand backward steps $[k_1 = (68.8 \pm 0.6)\text{s}^{-1}$ and $k_2 = (5.77 \pm 0.02)\text{s}^{-1}]$. (d) Inchworm backward steps $[k_1 = (67.5 \pm 0.6)\text{s}^{-1}$ and $k_2 = (5.76 \pm 0.02)\text{s}^{-1}]$.

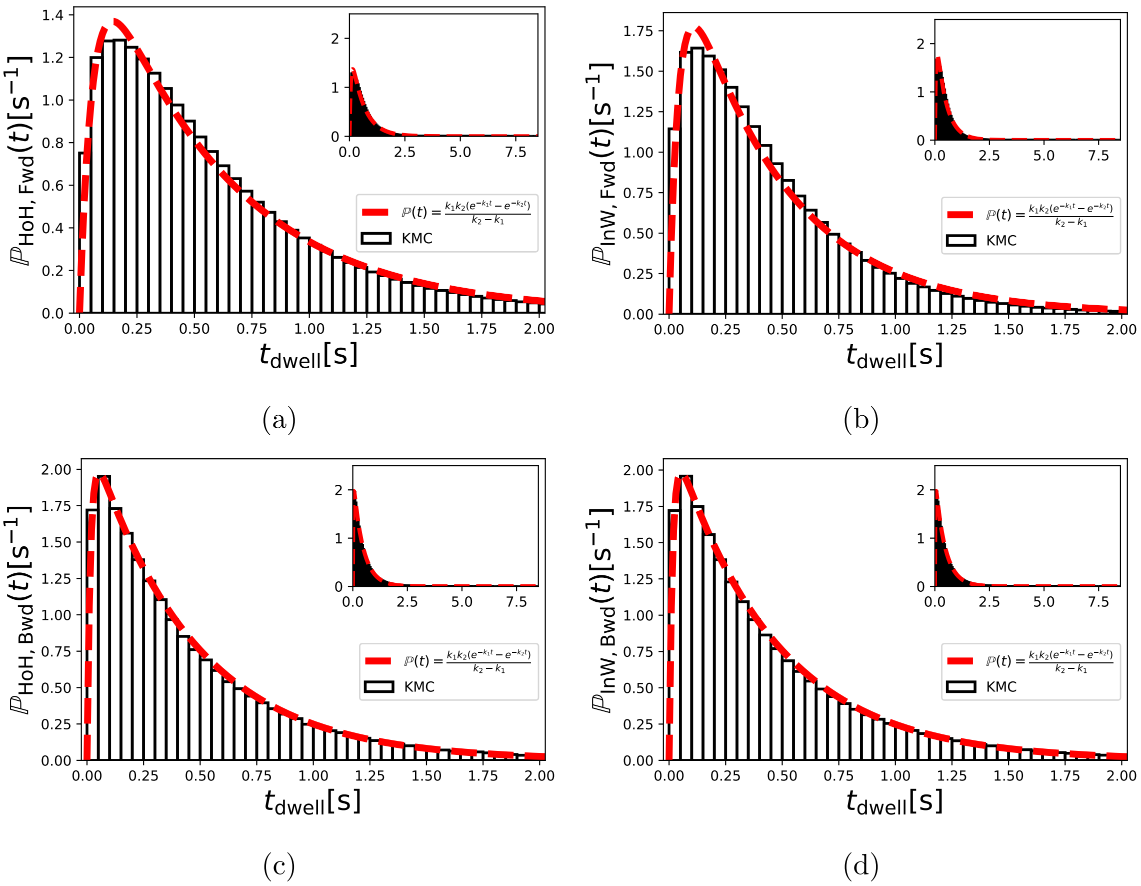


Figure S8: Dwell-time distributions at [ATP] = 100 $\mu$M. Histograms, in black, are from KMC simulations. For details, see caption of Fig. S7. From the fitting, the rates associated with hand-over-hand and inchworm forward steps are, (a) $k_1 = (17.7 \pm 0.4)\mathrm{s}^{-1}$ and $k_2 = (1.77 \pm 0.01)\mathrm{s}^{-1}$, and (b) $k_1 = (23.5 \pm 0.7)\mathrm{s}^{-1}$ and $k_2 = (2.28 \pm 0.02)\mathrm{s}^{-1}$, respectively. For backward steps we obtained, (c) $k_1 = (65.6 \pm 0.9)\mathrm{s}^{-1}$ and $k_2 = (2.239 \pm 0.006)\mathrm{s}^{-1}$, and (d) $k_1 = (65.7 \pm 0.8)\mathrm{s}^{-1}$ and $k_2 = (2.232 \pm 0.006)\mathrm{s}^{-1}$, for hand-over-hand and inchworm steps, respectively.

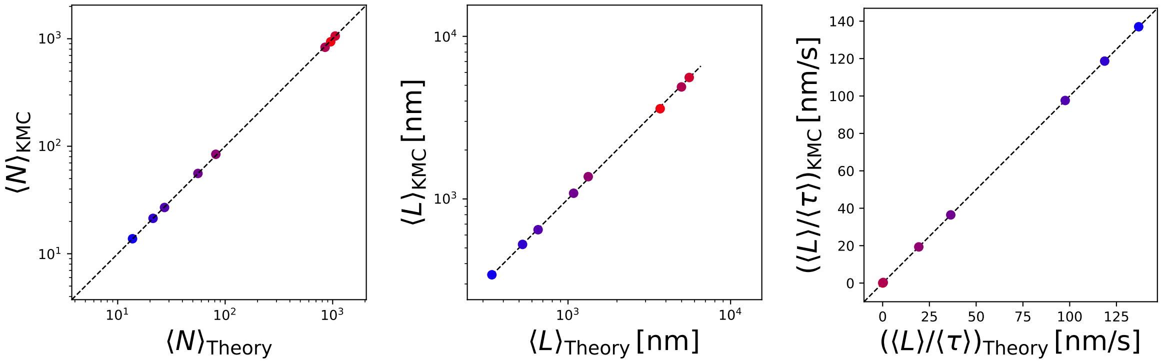


(a) (b) (c)

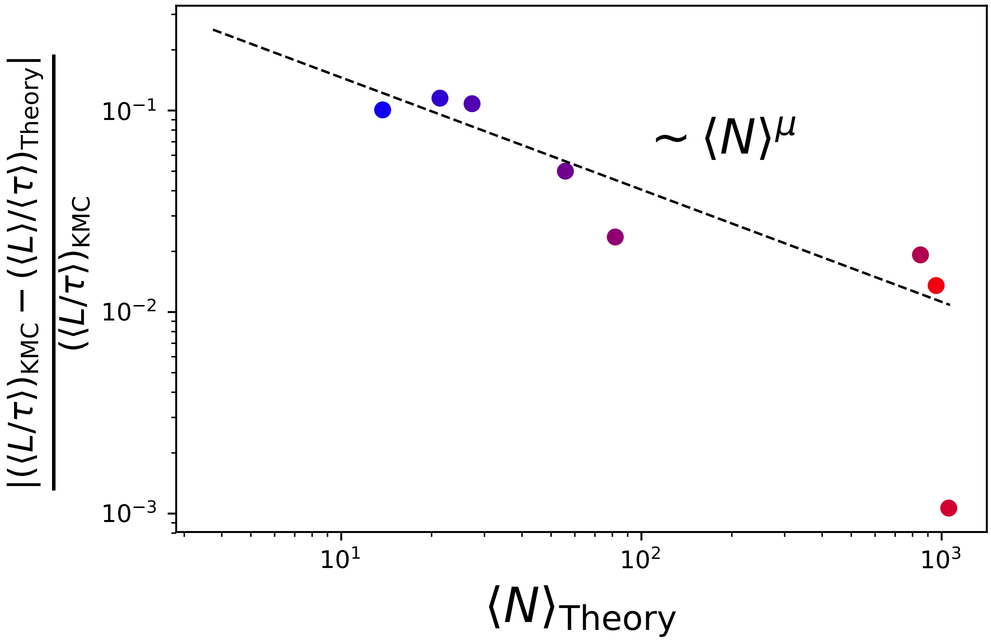


(d)

Figure S9: Comparison between two definitions of velocity $\langle L \rangle / \langle \tau \rangle$ and $\langle L/\tau \rangle$. The top three panels (a-c) show that the results from KMC simulations (y-axis) and the theory (x-axis) are essentially identical. The black line shows the function $y = x$. In panel (d) we show the relative deviation of $\langle L \rangle / \langle \tau \rangle$ computed with the the theory discussed in this paper and $\langle L/\tau \rangle$ obtained from KMC simulations as a function of the number of steps taken by the motor. The black line is an empirical fit to the power law $aN^{\mu}$, with $\mu = -0.56 \pm 0.15$ ($a$ and $\mu$ were obtained via fit). The points are computed at [ATP] of $0.1\,\mu$M, $0.5\,\mu$M, $1\,\mu$M, $50\,\mu$M, $100\,\mu$M, $500\,\mu$M, 1 mM, and 5 mM. The color of the point changes from red (low ATP concentration) to blue (high ATP concentration). The KMC simulations are averaged over an ATP-dependent number of trajectories: 1000 for [ATP] $= 0.1\,\mu$M, $0.5\,\mu$M, 5000 for $1\,\mu$M, and $50\,\mu$M, 10000 if [ATP] $= 100\,\mu$M, and $500\,\mu$M, and 100000 at the largest ATP concentrations ([ATP] $= 1$ mM, and 5 mM).

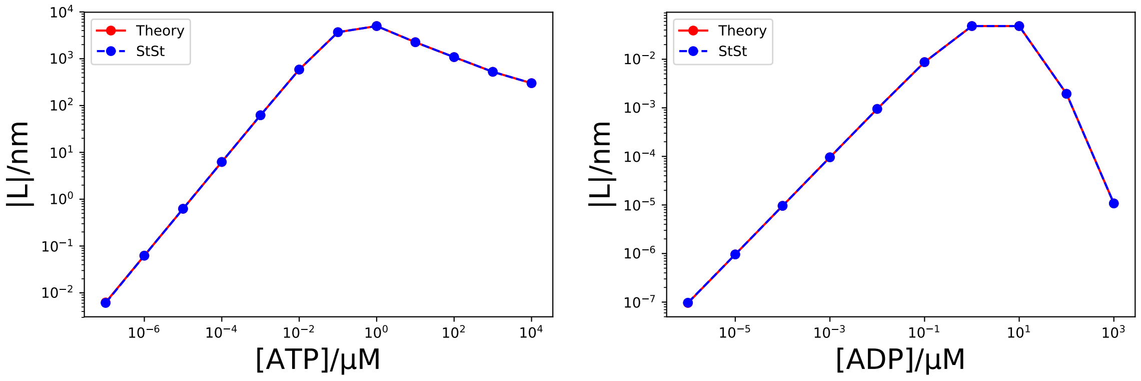


(a) (b)

Figure S10: Run length as a function of ATP and ADP. (a) Run length as a function of ATP with [ADP] = 0. (b) Dependence of the run length on ADP concentration in the absence of ATP. The full line connects points obtained using the equations described in the main text. The dashed line refers to calculations using the steady-state approach mentioned in the Discussion of the main text.
